# Insulin resistance compromises midbrain organoid neural activity and metabolic efficiency predisposing to Parkinson’s disease pathology

**DOI:** 10.1101/2024.05.03.592331

**Authors:** Alise Zagare, Janis Kurlovics, Catarina Almeida, Daniele Ferrante, Daniela Frangenberg, Laura Neises, Armelle Vitali, Gemma Gomez-Giro, Christian Jäger, Paul Antony, Rashi Halder, Rejko Krüger, Enrico Glaab, Johannes Meiser, Egils Stalidzans, Giuseppe Arena, Jens C Schwamborn

## Abstract

Growing evidence indicates that Type 2 Diabetes (T2D) is associated with an increased risk of developing Parkinson’s disease through shared disease mechanisms. Studies show that insulin resistance, which is the driving pathophysiological mechanism of T2D plays a major role in neurodegeneration by impairing neuronal functionality, metabolism, and survival. To investigate insulin resistance caused pathological changes in the human midbrain, which could predispose a healthy midbrain to PD development, we exposed iPSC-derived human midbrain organoids from healthy individuals to either high insulin concentrations, promoting insulin resistance, or to more physiological insulin concentrations restoring insulin signalling function. We combined experimental methods with metabolic modelling to identify the most insulin resistance-dependent pathogenic processes. We demonstrate that insulin resistance compromises organoid metabolic efficiency, leading to increased levels of oxidative stress. Additionally, insulin-resistant midbrain organoids showed decreased neural activity and reduced amount of dopaminergic neurons, highlighting insulin resistance as a significant target in PD prevention.

## Introduction

Parkinson’s disease (PD) is the second most frequent neurodegenerative disease, which affects the motor and cognitive function of patients due to the progressive degeneration of dopaminergic neurons in the *substantia nigra* region of the midbrain (1, 2). Most PD cases are idiopathic with an unknown genetic cause. Along with age as the primary risk factor, also environment, genetic risk factors and comorbidities contribute to the development of idiopathic PD (2). Recent epidemiological studies and meta-analyses show that Type 2 Diabetes (T2D) is associated with a 30 to 40% increased risk of developing PD (3–5). It has also been shown that patients affected by PD and T2D comorbidity, suffer from more severe motor symptoms and cognitive decline (6–9). Moreover, it has been suggested that T2D and PD share underlying pathophysiological mechanisms (5, 10, 11). Along with the dysregulated energy metabolism, aberrant protein aggregation, accumulation of reactive oxygen species and inflammation, insulin resistance seems to play a pivotal role in converging most of these common mechanisms. Insulin receptor presence in the basal ganglia and *substantia nigra* of healthy brains, and their loss in PD patient brains, suggests the particular importance of insulin in the regulation of dopaminergic neuron survival, as well as in the maintenance of their functional and metabolic homeostasis (12, 13). In addition, there is evidence showing that insulin resistance can occur in PD patient brains independently of T2D, highlighting insulin resistance as a potential factor contributing to PD onset and progression (13, 14).

In this study, we aimed to evaluate insulin resistance-mediated effects on the healthy midbrain to investigate through which molecular mechanisms insulin resistance might favour PD onset and progression. In order to address this question, we used induced pluripotent stem cell (iPSC)-derived human midbrain organoids from healthy donors and exposed them to a high insulin concentration, which mimics hyperinsulinemia and promotes insulin resistance, or to a more physiological insulin concentration to retain cells in an insulin-sensitive state. Using a combination of computational and experimental methods we predicted and validated insulin resistance-dependent metabolic alterations and demonstrated that insulin-resistant midbrain organoids experience functional decline in terms of electrophysiological activity. Further, they produce less dopamine and have a reduced amount of dopaminergic neurons. Overall these results support the notion that insulin resistance is a considerable risk factor for PD and that it can be modeled *in vitro*

## Results

### Modification of cell culture media activates insulin signalling

Midbrain organoids were generated from three healthy individual iPSC-derived neuroepithelial stem cell lines (NESCs) (Table S1) following a previously published protocol (16, 18, 19). Before organoid generation, the quality of all NESC lines used in the study was confirmed by evaluating the expression of the stem cell marker SOX2, the neural progenitor marker Nestin and the neuroectoderm fate marker PAX6 (Figure S1). Midbrain organoids were cultured in two culture conditions that vary by the concentration of insulin (Figure 1A). First, we determined the insulin concentration in standard basal N2B27 media supplemented with insulin-containing neurotrophic supplements B27 and N2 used for the maintenance of NESCs and midbrain organoids. The insulin concentration in the standard basal media was 393.9 nM, which strongly exceeds the physiological insulin concentration in the central nervous system, reported to be below 0.1nM (measured in cerebrospinal fluid) (20, 21). We hypothesised that this supraphysiological insulin concentration in the standard media might cause insulin resistance (IR) in the organoid culture. We also measured insulin concentration in the supernatant of organoids cultured for three days in the standard media with the high insulin concentration and saw that the insulin concentration remained between 352 nM and 374 nM (Figure 1B). Next, to reduce insulin concentration in the media towards a more physiological one, we prepared a self-made N2 supplement following commercial N2 media formulation, only excluding insulin (see material and methods). Insulin concentration in the N2B27 medium containing self-made N2 was 171.2 nM which is 2.3 times less than in the standard media. After three days of organoid culturing in this reduced insulin media, insulin concentration decreased even more, and was between 103.7 nM and 127.4 nM, suggesting increased usage of insulin by cells (Figure 1B). Further, we confirmed that self-made N2 supplement did not affect organoid development nor viability (Figure S2) and did not change cellular composition within organoids (Figure S3). Since dopaminergic neurons are the most relevant cell type for PD, we additionally validated the self-made N2 supplement on iPSC-derived dopaminergic neuron cultures. Self-made N2 media did not cause any changes in dopaminergic neuron differentiation capacity, determined by the similar levels of TH and TUJ1 markers in dopaminergic neurons cultured in standard and self-made N2 supplemented media (Figure S4A). To further confirm that a 2.3 times reduction of the insulin concentration is sufficient to change the activity of insulin signalling, we cultured midbrain organoids in these two insulin conditions from the start of organoid differentiation – day two of organoid culture and maintained until the sample collection – day 30 or day 60 of organoid culture (Figure 1A).

**Figure 1.**
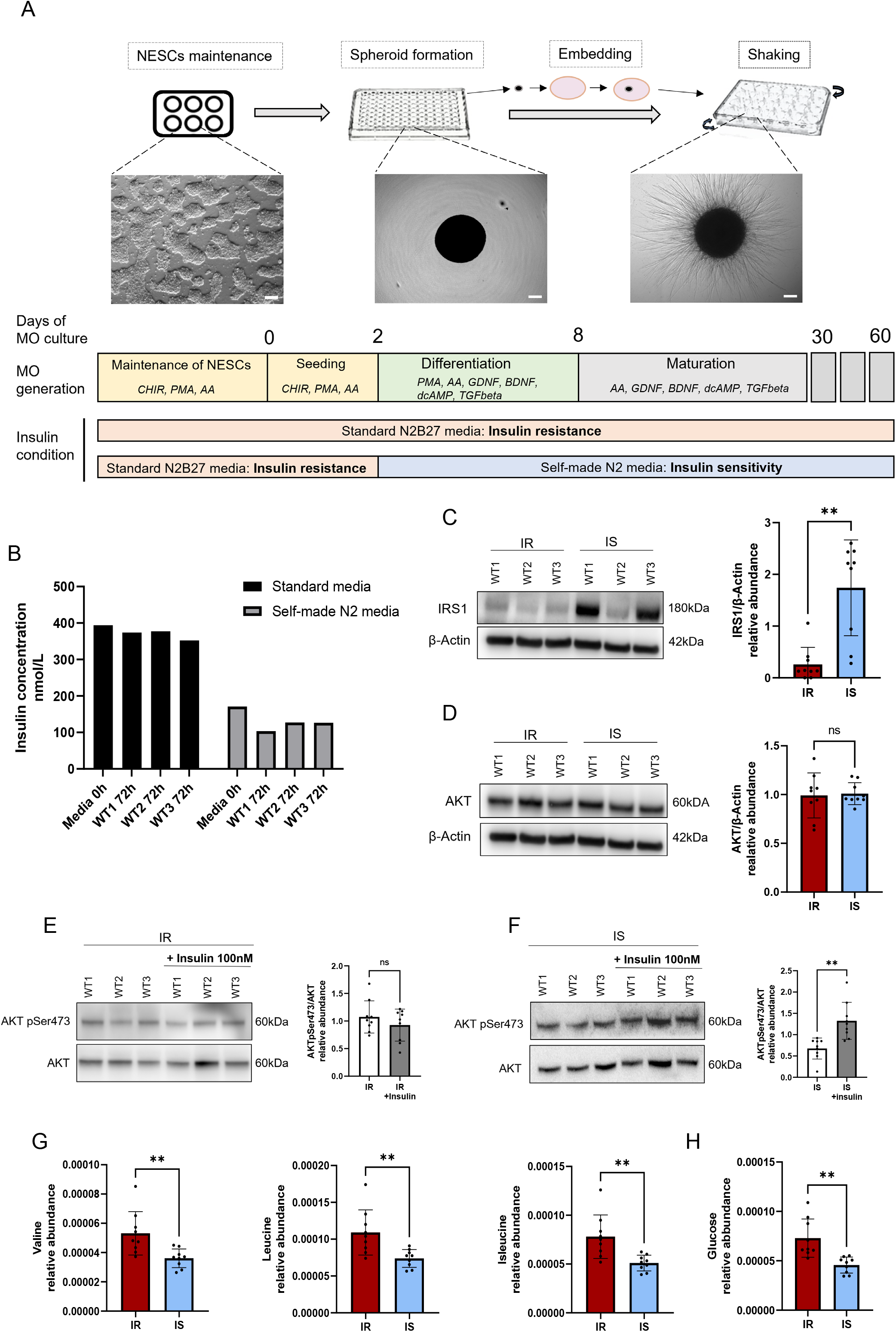
Modification of cell culture media activates insulin signalling. (A) Representative image of midbrain organoid generation and cell culture media conditions. (B) Insulin concentration in standard media and self-made N2 media measured in fresh media (0h) and after three days of incubation with cells (72h). Scale bars 200µm. (C) Representative image of WB and quantification of IRS1 levels relative to the housekeeping β-actin. (D) Representative image of WB and quantification of AKT1 levels relative to the housekeeping β-actin. (E) Representative image of WB and quantification of AKT1 pSer73 levels after acute stimulation with 100nM of insulin in IR samples, relative to the total AKT levels. (F) Representative image of WB and quantification of AKT1 pSer73 after acute stimulation with 100nM of insulin in IS samples, relative to the total AKT levels. For panels C-F statistical significance was tested with two-tailed, match-pairs signed ranked Wilcoxon test (*P*<0.05 *, *P* <0.01 **, *P* <0.001 ***). Error bars represent mean + SD. (G) Relative abundance of branched chain amino acids – Valine, Leucine, and Isoleucine, detected by LC-MS in midbrain organoid supernatant after 72h incubation with organoid. (H) Relative abundance of glucose, detected by LC-MS in midbrain organoid supernatant after 72h incubation with organoid. For panels g-h statistical significance was determined with a two-tailed, paired t-test (*P*<0.05 *, *P* <0.01 **, *P* <0.001 ***). Error bars represent mean + SD.

We were able to demonstrate a significant reduction of insulin receptor substrate (IRS1) upstream insulin signalling in organoid samples cultured in the high insulin concentration standard media, confirming insulin signalling dysfunction in these organoids (Figure 1C). On the contrary, IRS1 was significantly more abundant in the organoid samples cultured in reduced insulin concentration media, suggesting that this condition promotes cellular insulin sensitivity (IS). Additionally, we looked at the abundance of the central protein of insulin signalling – AKT, which was not changed between organoids cultured in the two insulin conditions (Figure 1D). We triggered acute insulin signalling activation by organoid stimulation with 100nM insulin for two hours and observed that in response to insulin, there was an increase of AKT phosphorylation at Ser473 residue in IS organoids, while IR organoids did not show any change in AKT phosphorylation (Figure 1E-F), further demonstrating loss of insulin sensitivity. Similarly, the acute stimulation of only dopaminergic neurons with increasing concentrations of insulin was able to activate the insulin signalling pathway already at the dose of 10nM, as indicated by the robust increase of AKT phosphorylation (both at Ser473 and Thr308 residues) in dopaminergic neurons cultured in self-made N2-based media. In contrast, even the highest dose of insulin (200nM) was unable to increase phospho-AKT levels in dopaminergic neurons cultured in N2B27 media containing commercial N2, further supporting that insulin concentration change affects the insulin signalling also in dopaminergic neurons (Figure S4B).

To additionally confirm the IR and IS state of midbrain organoids cultured under different insulin concentration media, we measured levels of branched-chain amino acids which are considered biomarkers of insulin resistance and are found in higher levels in the plasma of individuals with changed metabolism and insulin resistance (22, 23). Accordingly, we saw their increased levels in IR organoid-conditioned media (Figure 1G). Furthermore, we observed that most of the metabolites present in the culture media were in significantly lower concentration in the IS organoid spent media after three days of incubation, indicating better metabolite uptake by IS organoids (Figure S5A). These metabolites included glucose, which was also found in significantly decreased levels in IS organoid spent media (Figure 1H). To verify increased glucose uptake by IS organoids, we treated dissociated organoids with a fluorescent glucose analogue 2-NBDG, to visualise glucose uptake by fluorescence-activated cell sorting (FACS) (Figure S5B). We were able to observe a significantly increased fluorescent signal for IS organoids after acute insulin stimulation, further supporting that they are more insulin-responsive than IR organoids. To confirm that glucose uptake in midbrain organoids can be insulin-dependent, we investigated the glucose transporter (GLUT) presence in midbrain organoids. Along with the insulin-independent GLUTs, such as neuron-specific GLUT3 and glia-specific GLUT1, we were able to detect insulin-dependent GLUT4, which supports insulin’s role in glucose uptake and regulation of metabolism in midbrain organoids (Figure S5C). Moreover, we observed that GLUT4 staining overlapped with the neuronal marker MAP2 (Figure S4D) and the glia marker GFAP (Figure S4E), suggesting GLUT4 expression in these cell types. Furthermore, we did not observe a significant difference in SOX2+ stem cells, MAP2+ neurons and the S100b+ glia population between IR and IS midbrain organoids at day 30 nor day 60 of organoid culture (Figure S3), hence we conclude that here presented metabolic differences are rather insulin concentration-dependent and not the consequence of changes in the cell type proportions within midbrain organoids.

### Insulin sensitivity increases dopaminergic neuron amount and midbrain organoid functionality

After we confirmed that a more physiological insulin concentration in the cell culture restores insulin signalling activity in the whole midbrain organoid as well as in the dopaminergic neurons, we wanted to assess whether insulin concentration affects dopaminergic neuron abundance. While at an earlier time point (day 30), there was no significant difference in the amount of TH-positive dopaminergic neurons between the two conditions, we observed a significantly higher number of dopaminergic neurons in IS midbrain organoids at day 60 of organoid culture (Figure 2A-B). Moreover, we confirmed that increased dopaminergic neuron amount in IS organoids is accompanied by a significant increase in dopamine production (Figure 2C). Considering the positive effect of insulin sensitivity on dopaminergic neuron levels and their ability to produce dopamine, we wanted to assess the role of insulin in the electrophysiological activity of the entire midbrain organoid. We measured spontaneous electrophysiological activity in the midbrain organoids using a multi-electrode array (MEA) system (Figure 2D) and observed that organoids cultured in IS conditions presented a significantly higher mean firing rate (Figure 2E). Furthermore, we confirmed the increased electrophysiological activity of IS organoids by recording local field potentials (LFP). Supporting the MEA data, IS organoids demonstrated a significantly higher number of spikes than IR organoids at both time points with an increased spike duration at day 60 of culture (Figure 2F-H). We were also able to detect bursts, which represent synchronized neuronal activity, occurring more often in IS organoids than in IR organoids again at both time points (Figure 2I-K), further confirming that insulin sensitivity promotes neuronal functionality.

**Figure 2.**
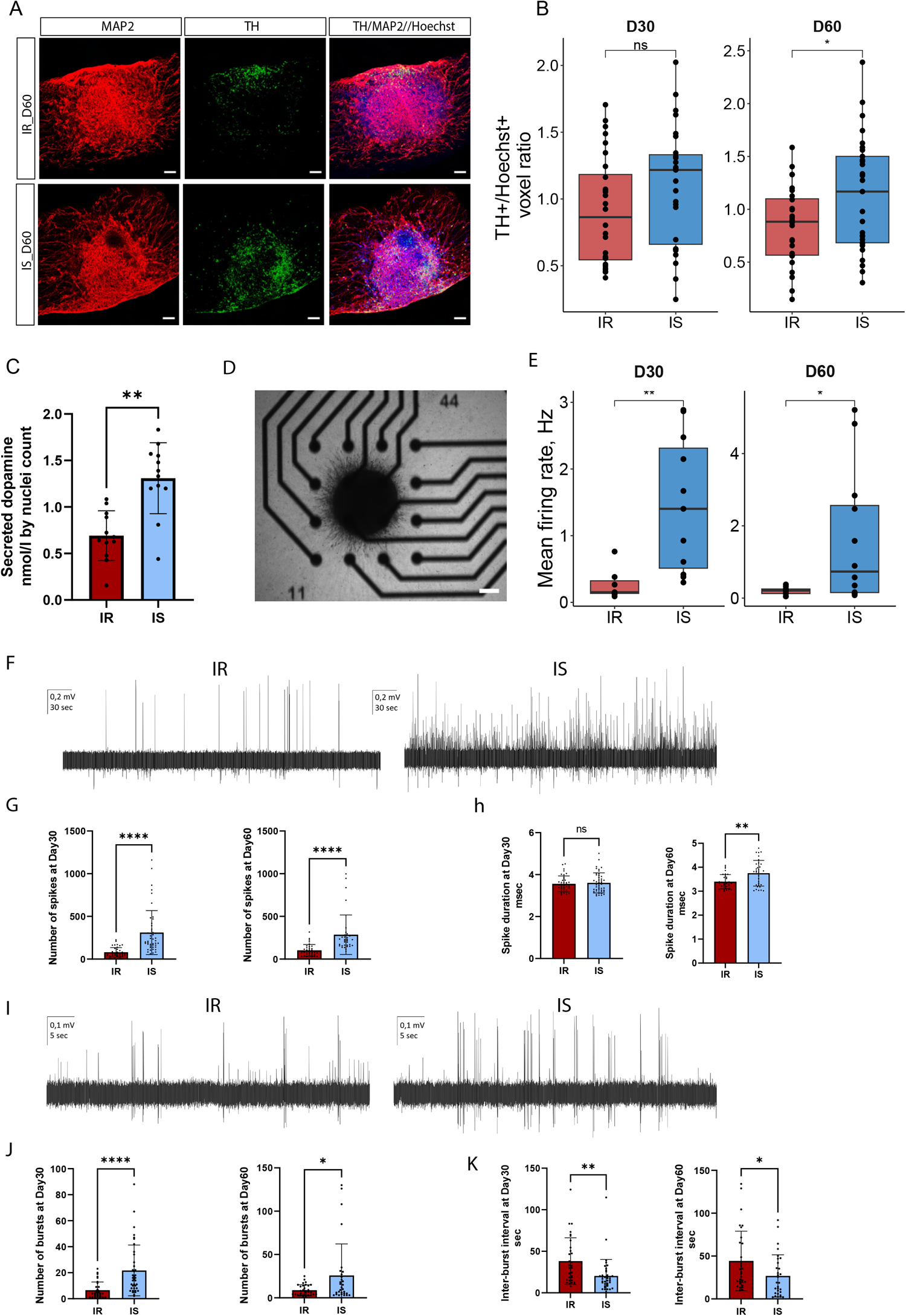
Insulin sensitivity increases dopaminergic neuron amount and midbrain organoid functionality. (A) Representative immunofluorescent images of TH and MAP2 staining in sections of 60 days old midbrain organoids. Scale bars 100µm. (B) Quantification of TH positive area normalised to the Hoechst area in immunofluorescent stainings of 30 and 60 days old midbrain organoids. (C) Extracellular dopamine levels measured in spent media of 60 days old midbrain organoids. (D) Representative image of organoid placed on electrodes in a well of the multielectrode plate. Scale bar 200µm. (E) Mean firing rate of 30 and 60 days old midbrain organoids Recording time-5min. For panels B-E statistical significance was tested with a two-tailed Wilcoxon test (*P*<0.05 *, *P* <0.01 **, *P* <0.001 ***). (F) Representative image of local field potential recording of spontaneous firing in midbrain organoids at day 30. (G) Quantification of spikes per time of recording (5min) for day 30 and day 60 old organoids. (H) Quantification of spikes duration in seconds for day 30 and day 60 old organoids. (I) Representative image of local field potential recording of burst firing in midbrain organoids at day 60. (J) Quantification of bursts per time of recording (5min) for day 30 and day 60 old organoids. (K) Quantification of inter-burst interval in seconds for day 30 and day 60 old organoids. For panels G-K statistical significance was tested with Mann–Whitney t-test (*P*<0.05 *, *P* <0.01 **, *P* <0.001 ***, *P* <0.0001 ****). Error bars represent mean + SD.

### Metabolic modelling predicts altered lipid metabolism and decreased glycolysis efficiency under insulin resistance

To further investigate metabolic alterations linked to insulin resistance, we performed metabolic modelling integrating RNA sequencing data of IR and IS midbrain organoids into the Recon3 human metabolic network (24). Gene expression was integrated using the rFASTCORMICS pipeline (25) which builds metabolic models based on gene discretization (Figure S6A-B). We generated IR and IS models from pooled transcriptomic data between three cell lines cultured in the respective conditions. IR and IS model composition was not substantially different (Figure S6C). The IR model included 1446 genes, 3620 metabolic reactions and 2632 unique metabolites. While the IS model included 1430 genes, 3470 metabolic reactions and 2521 unique metabolites. Both models were primarily constrained by the N2B27 basis media composition, allowing uptake of only those metabolites that are present in the media. To capture differences in metabolism more accurately, we integrated major metabolic constraints such as glucose uptake and lactate production. We also added biomass constraint, as organoid growth between day 16 and day 30 of culture to account for biomass change per unit of time, allowing more precise estimation of required metabolic rates of reactions present in the models.

By model composition analysis we identified the most different pathways by reaction composition between the two models (Figure 3A). The top metabolic pathways included several transport subsystems between cellular compartments and lipid metabolism-associated metabolic subsystems. To confirm that altered lipid metabolism is driven by insulin resistance, we validated our modelling results by lipidomic analysis, which detected an abundance of 18 different lipid classes in midbrain organoid samples. The lipid profile separated IR and IS samples in a principal component analysis (PCA) across the first principal component (11.6%), although the variance between the samples across the second component was higher (63,5%), demonstrating biological variability between the cell lines. We found that seven lipid classes were significantly differentially abundant between IR and IS samples (Figure S7), the most significantly dysregulated class and the major contributor to the IR sample separation were cholesterol esters (Figure 3C-D). Consistent with the *in silico* predicted alteration in glycerophospholipid metabolism, lipidomics analysis revealed that phosphatidylcholines, which are one of the most abundant cell membrane phospholipid classes, were significantly increased in IS samples (Figure 3E). Additionally, modelling predicted changes in arachidonic acid metabolism, therefore, we investigated lipid species across all measured lipid classes that contained arachidonic acid (C20:4) and its immediate precursor of eicosanoid family – eicosatrienoic acid (C20:3). We found 10 significantly differentially abundant species containing a 20-carbon backbone across six lipid classes, all being intermediates of glycerophospholipid metabolism, namely – triacylglycerides (TG), phosphatidylcholine (PC), 1-alkyl,2-acylphosphatidylethanolamines (PC-O), 1-alkenyl,2-acylphosphatidylethanolamines (PC-P), lysophosphatidylcholine (LPC), lysophosphatidylethanolamine (LPE) (Figure S8). Here the best separation between the IR and IS samples was achieved by major lipid components of the cellular membrane, including PC-Ps, PC-Os and the phosphatidylcholine*s* derivatives LPCs (Figure 3F). All together these results indicate that insulin resistance is inducing lipid changes that particularly affect cellular membrane and cholesterol metabolism.

**Figure 3.**
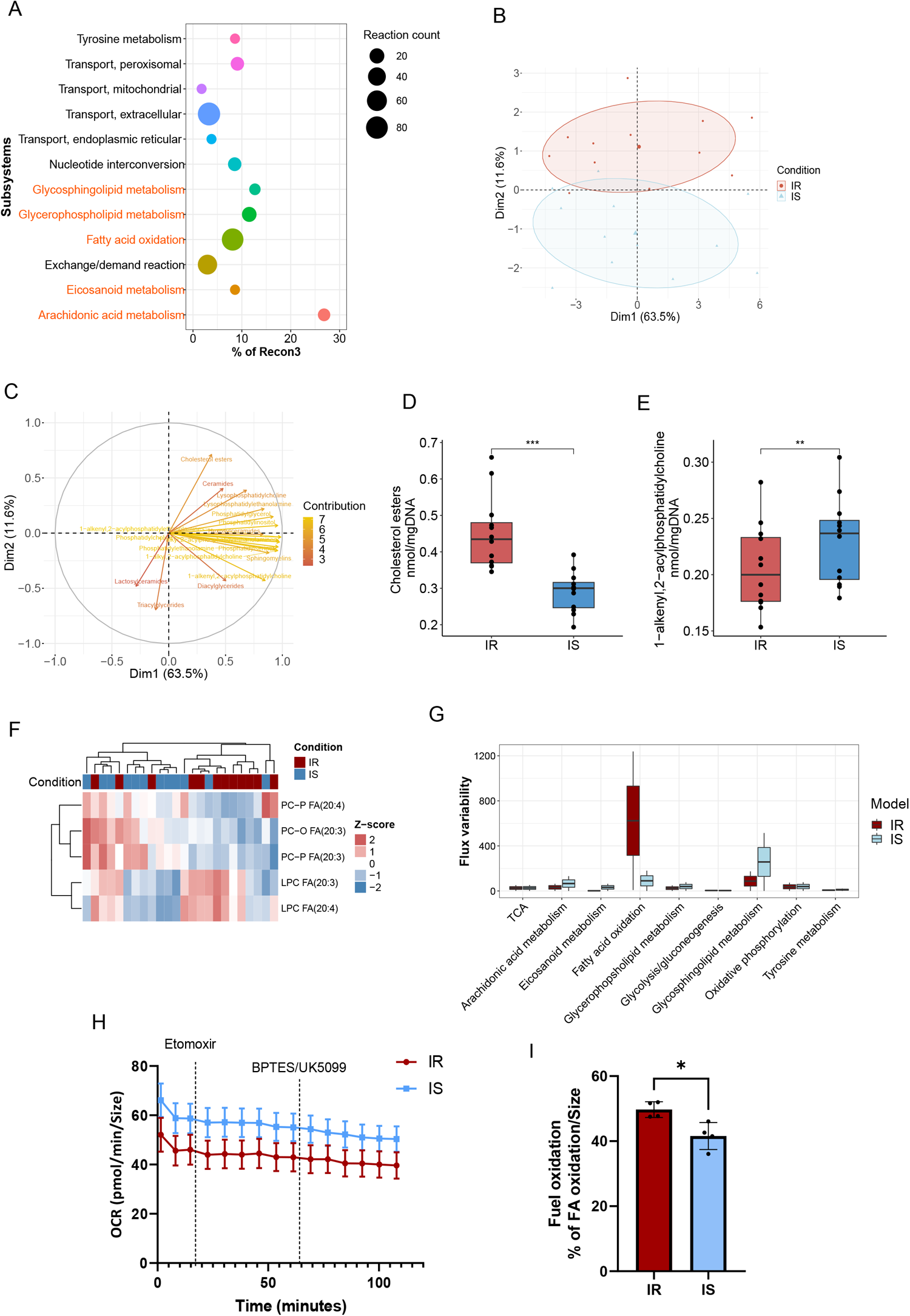
Insulin resistance-mediated lipid alterations. (A) Prediction of the most dysregulated metabolic pathways in the IR model based on model composition analysis. The size of a dot represents the number of reactions different between models. The location on the x-axis represents the percentage of reactions different from the total reaction number per subsystem in Recon3. Highlighted subsystems belong to lipid metabolism. (B) PCA scores plot of lipidomics analysis, demonstrating the percentage of variation between IR and IS sample lipidomic profiles, explained by the first and second components. (C) Correlation plot, demonstrating feature (individual lipid class) correlation across the samples and contribution to the variance between the conditions. (D) Quantification of Cholesterol esters. Concentration normalised to the mgDNA. (E) Quantification of 1-alkenyl,2-acylphospatidylcholine. Concentration normalised to the mgDNA. For panels D-E statistical significance was tested with a two-tailed, match-pairs signed ranked Wilcoxon test (*P*<0.05 *, *P* <0.01 **, *P* <0.001 ***). (F) Unsupervised clustering of the most significantly differentially abundant lipid species derived from 20-carbon arachidonic or eicosanoid acid. (G) Prediction of metabolic pathway performance by flux variability analysis (FVA). (H) Fatty acid dependency measurements obtained using Seahorse assay. (I) Quantification of the percentage of fatty acid oxidation from the total fuel oxidation. Results normalised to the size of organoids. Statistical significance was tested using paired two-tailed t-test (*P*<0.05 *). Error bars represent mean + SD.

Next, we carried out an *in silico* flux variability analysis (FVA) to predict the differences in metabolic performance of the subsystems of our interest. We included lipid subsystems with high differences in the composition between the models (Figure 3A), and central energy metabolism pathways – glycolysis, TCA and OXPHOS (Figure 3G). We observed higher flux flexibility in glycosphingolipid metabolism in the IS model compared to the IR model, suggesting that insulin resistance might impair glycosphingolipid metabolism. In contrast, the IR model showed much higher flexibility in fatty acid oxidation than the IS model, suggesting that fatty acid oxidation might contribute to energy generation in insulin-resistant conditions. We wanted to confirm this prediction by Seahorse metabolic assay, which allows us to evaluate organoid metabolic dependency on different carbon sources for energy generation. Indeed, we observed that IR organoids are about 10% more dependent on fatty acids than IS organoids (Figure 4H-I). We further performed *in silico* perturbations to determine the importance of glucose uptake and mitochondrial performance for the generation of ATP (Figure 4A). ATP levels were predicted to be slightly higher in the IS model compared to the IR model. We confirmed this by measuring intracellular ATP levels in 30 and 60 days old organoids, demonstrating significantly higher levels of ATP in IS samples at both time points (Figure 4B-C). There was no predicted difference in ATP generation by models after *in silico* blocking mitochondrial complexes I and V. However, the prediction showed that the IS model is unable to generate ATP in case of glucose uptake obstruction (Figure 4A). This suggested that the IS model strongly relies on glucose to produce ATP. At the same time the IR model showed an irrelevant change in ATP levels in case of blocking glucose uptake, implying possible usage of other carbon sources for energy generation under insulin resistance conditions. Additionally, we performed glucose C13 tracing to confirm increased glycolytic flux in IS samples. Indeed, we saw a significant increase in the relative labelled fraction of glycolytic intermediates directly derived from labelled glucose – 3-phosphoglycerate (3PG), phosphoenolpyruvate (PEP), pyruvate and lactate in IS samples (Figure 4D). However, tracing of labelled glucose did not reveal a labelling difference in TCA intermediates, suggesting no strong functional differences in mitochondrial metabolism between IR and IS midbrain organoids. In addition, we observed higher relative labelled fraction of Ribose 5-phosphate and Serine in the IS samples, compared to IR samples, pointing towards increased glucose derived carbon distribution to the pentose-phosphate pathway and *de novo* Serine synthesis respectively (Figure 4E-F). Further, we anticipated that altered metabolism in IR organoids might cause increased oxidative stress which plays an important role in the pathophysiology of PD. Accordingly, we detected significantly lower levels of NAD+/NADH ratio in IR samples, indicating an altered redox state of IR organoids. We also observed increased cytosolic ROS levels (Figure 4H-I), further confirming metabolic and redox disbalance in midbrain organoids driven by insulin resistance.

**Figure 4.**
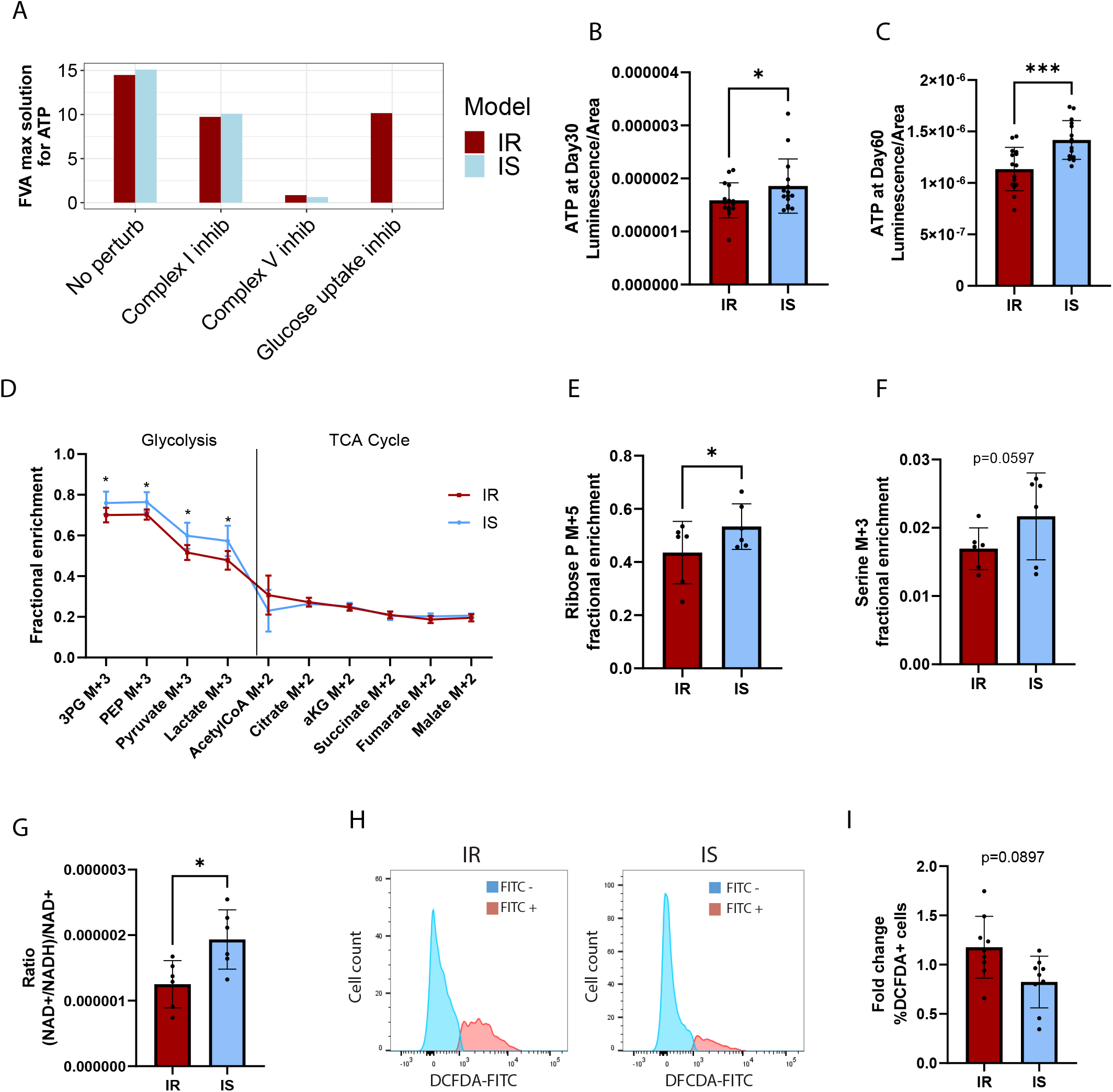
Insulin resistance compromises ATP levels and glycolysis efficiency. (A) Prediction of ATP production by IR and IS model following *in silico* inhibition of mitochondrial complexes present in the model and glucose uptake inhibition. (B) Intracellular ATP measurement for organoids at day 30 of organoid culture. (C) Intracellular ATP measurement for organoids at day 60 of organoid culture. (D) Fractional abundance of glycolysis and TCA metabolites derived from U^13^C labelled glucose. For panels b-d statistical significance was tested with a two-tailed, match-pairs signed ranked Wilcoxon test (*P*<0.05 *, *P* <0.01 **, *P* <0.001 ***). Error bars represent mean + SD (E) Quantification of Serine derived from U^13^C labelled glucose. (F) Quantification of Ribose 5-phosphate derived from U^13^C labelled glucose. (G) The ratio of total and reduced NAD normalised to the total NAD levels. (H) Flow cytometry representative histograms, showing DCFDA positive (FITC-A+) fraction of cells in IR and IS condition. (I) Quantification of live cells positive to DCFDA (FITC-A) fluorescent signal. For panels E-I statistical significance was tested with a two-tailed, paired t-test (*P*<0.05 *, *P* <0.01 **, *P* <0.001 ***, *P* <0.0001 ****). Error bars represent mean + SD

## Discussion

In this study, we aimed to understand the molecular mechanisms through which T2D-associated insulin resistance might contribute to onset and progression of PD. With a global prevalence of over 10% and a tendency to increase (26, 27), T2D has become a considerable risk factor for many diseases, including neurodegenerative disorders. Thus, there is a need to better understand T2D contribution to the onset and progression of these diseases. Moreover, PD is currently the fastest-growing neurodegenerative condition with still unclear molecular causes of neuronal death in the vast majority of cases (28). Identification of risk factors might open new preventative strategies, halting the rising number of PD cases in future. Here we show that insulin resistance, a hallmark of T2D, predisposes a healthy midbrain to PD-associated neurodegeneration by considerably altering metabolism and compromising the amount of dopaminergic neurons, which is the main affected neuronal type in PD.

It has been previously reported that not only T2D but even pre-diabetes with insulin resistance as a common factor, promotes cognitive decline in PD patients (7, 29). We were able to demonstrate that under insulin resistance midbrain organoids display reduced spontaneous activity in the form of action potentials and bursts, suggesting decreased neuronal functionality, which could lead to cognitive impairment. Brain energetics is crucial to coordinate electrophysiological activity (30, 31) and since insulin signalling is a major regulator of metabolic processes, it is expected that insulin resistance would affect neuronal signal transmission. Although brain insulin sensitivity is controversial, mouse studies have shown that in some brain regions for rapid glucose uptake neurons are equipped with insulin-dependent glucose transporters in the proximity of synapses (32, 33), supporting insulin’s role in controlling brain energy metabolism. Accordingly, we were able to show the presence of insulin-dependent GLUT4 along with the insulin-independent GLUT1 and GLUT3 transporters in midbrain organoids, suggesting at least partial insulin-dependent glucose uptake in the human midbrain. This also means that in the case of insulin resistance, neuronal functionality might be compromised due to insufficient glucose uptake for fast energy requirements. A reduced glycolysis rate in IR organoids we also confirmed with glucose C13 tracing. However, we did not observe any changes in levels of TCA intermediates derived from labelled glucose. This suggests that all glucose uptaken by cells in IR midbrain organoids is completely metabolized in the mitochondria for energy generation, probably to restore reduced levels of ATP. Moreover, we observed a slight but significant increase in fatty acid oxidation by IR organoids, which might present cell attempts to ensure sufficient energy levels in case of glucose unavailability. However, fatty acid oxidation is a considerable source of superoxide, which with time can exhaust anti-oxidative mechanisms and lead to cell damage (34). At the same time, increased glucose uptake by IS organoids allows glucose usage for other metabolic pathways, such as *de novo* serine synthesis and nucleotide synthesis via the pentose-phosphate pathway. Both pathways are important for healthy brain function, playing a role in cell survival, neurotransmission, regulation of redox state, and lipid and protein synthesis (35, 36).

In addition, to the altered rate of fatty acid oxidation, we observed significant changes in abundance levels of several important lipid classes in IR midbrain organoids. Previous studies have reported elevated cholesterol ester levels associated with neurodegenerative diseases (37–39). In line, we detected increased levels of cholesterol esters in IR organoids, suggesting that they undergo molecular changes linked to neurodegeneration. In addition, we demonstrated that most of the cellular membrane lipid classes containing arachidonic acid are differentially abundant between IR and IS midbrain organoids. Arachidonic acid is one the most abundant fatty acids in the brain and it has been associated with the prevention of α-Synuclein protein aggregation (40) and neuroprotection in animal models (41). Moreover, it has been reported as a potential biomarker in PD (42), suggesting its relevance in disease progression. Additionally, one of the most abundant cell membrane phospholipids – phosphatidylcholines were found to be significantly reduced in IR organoids, suggesting insulin resistance effect on cellular membrane integrity. Interestingly, metabolic modelling predicted changes in glycosphingolipid metabolism, which might play a role in phenotype acceleration in PD cases caused by a mutation in the GBA1 gene. Heterozygous mutation in the GBA1 gene has been identified in 5-15% of all monogenic PD cases and is a major genetic risk factor for PD, associated with the accumulation of ceramides and glycosphingolipids - hexocylceramides, promoting PD pathophysiology (43–45). Thus, altered glycosphingolipid metabolism might be an important link between the GBA1-associated PD and insulin resistance.

Importantly, we observed that midbrain organoids cultured under insulin-resistant conditions have significantly reduced amounts of dopaminergic neurons. Since the dopaminergic neurons are sensitive to metabolic changes due to their high energy needs, their loss in insulin resistance might be primarily explained by insulin resistance-caused metabolic alterations. Dopaminergic neurons are the most affected cellular population in PD, thus their loss in IR midbrain organoids, strongly indicates that insulin resistance predisposes healthy midbrain to PD development and might be a potential target in the prevention of neurodegeneration.

## Data and code availability

All data supporting the conclusions of this study are publicly available at: DOI: https://doi.org/10.17881/ztcz-2197

RNA sequencing data is available on Gene Expression Omnibus (GEO) under the accession code GSE237116.

Matlab and R scripts for data analysis and modelling are available on GitHub at: https://github.com/LCSB-DVB

## Materials and Methods

### IPSC-derived dopaminergic neuron differentiation and characterisation

iPSC from a healthy donor were first converted into neuroepithelial stem cells (NESCs) based on the protocol previously described (28). In brief, the N2B27 medium was used as the basis media for cell culture. N2B27 consists of DMEM-F12 (Thermo Fisher Scientific, cat.no 21331046) and Neurobasal (Thermo Fisher Scientific, cat.no.10888022) 50:50 and is supplemented with 1:200 N2 supplement (Thermo Fisher Scientific cat.no. 17502001), 1:100 B27 supplement w/o Vitamin A (Life Technologies, cat.no. 12587001), 1 % GlutaMAX (Thermo Fisher Scientific, cat.no. 35050061) and 1 % penicillin/streptomycin (Thermo Fisher Scientific, cat.no. 15140122). N2B27 base media was supplemented with 3 µM CHIR-99021 (Axon Medchem, cat.no. CT 99021), 0.75 µM purmorphamine (Enzo Life Science, cat.no. ALX-420-045) and 150 µM ascorbic acid (Sigma, cat.no. A4544). For the generation of midbrain-specific dopaminergic neurons, NESCs were seeded into Geltrex-coated wells in a differentiation medium consisting of N2B27 containing 1 µM PMA, 200 µM ascorbic acid, 10 ng/ml BDNF, 10 ng/ml GDNF, 1 ng/ml TGF-b3 and 500 µM dbcAMP. After six days, differentiating NESCs were shifted to the same medium but without PMA (replaced every two days) and kept until day 30.

The expression of neuronal differentiation markers was assessed by flow cytometry. Briefly, cells were detached with Accutase (Sigma, cat.no. A6964) and then fixed in 4% PFA for 15 minutes. After three washes in FACS buffer (PBS + 5% BSA + 0.1% Sodium Azide), cells were centrifuged at 700g for 5 minutes and the resulting cell pellet resuspended in Saponin buffer (PBS + 1% BSA + 0.05% Saponin). After 30 minutes of incubation at 4°C, cells were pelleted again and then resuspended in Saponin buffer containing the TH (Millipore, cat.no. AB152) and Tuj1 (Biolegend, cat.no 801201) primary antibodies (1:100 dilution, 30 minutes at 4°C under shaking). Cells were washed three times in FACS buffer, and then incubated for 30 minutes with anti-mouse Alexa 488 (for TUJ1) and anti-rabbit Alexa 568 (for TH) secondary antibodies (at 4°C in the dark). After two washes in the FACS buffer, cells were pelleted and finally resuspended in PBS. Cells were analyzed with the BD LSRFortessa flow cytometer and the mean fluorescence intensity of each dye was assessed on at least 10,000 single cells using the FlowJo software (v.10.7.2.).

### Midbrain organoid culture

Cell lines used in this study are summarised in Table S1. As starting population for midbrain organoid generation, we used NESCs described in Reinhardt et al., 2013 (28). NESCs were cultured as described in paragraph *iPSC-derived dopaminergic neuron differentiation and characterisation*. Midbrain organoids were generated as described in Monzel et al.,2017 (29) and Nickels et al., 2020 (30). For organoid generation, at 80% of confluency, NESCs were detached using Accutase (Sigma, cat.no. A6964) and viable cells were counted using Trypan Blue. 9×10^5^ cells were transferred into 15ml of N2B27 maintenance media for seeding in 96-well ultra-low attachment plates (faCellitate, cat.no. F202003). In each well 150µl with 9000 cells were distributed. Plates were centrifuged at 300 g for one minute to facilitate colony formation at the bottom of the well. On the 2^nd^ day of organoid culture, media was changed to N2B27 pattering media, which is N2B27 base media supplemented with 10 ng/ml hBDNF (Peprotech, cat.no. 450-02-1mg), 10 ng/ml hGDNF (Peprotech, cat.no. 450-10-1mg), 500 µM dbcAMP (STEMCELL Technologies, cat.no.100-0244), 200 µM ascorbic acid (Sigma), 1 ng/ml TGF-β3 (Peprotech cat.no. 100-36E) and 1 µM purmorphamine (Enzo Life Science, cat.no ALX-420-045). Media was next changed on the 5^th^ day of organoid culture. On day 8 of organoid culture, organoids for high-content imaging were embedded in extracellular matrix-like Gelltrex (Thermo Fisher Scientific, cat.no. A1413302) droplets as described in Zagare & Gobin, et al., 2021 (31) to support the outgrowth of neurites. Embedded organoids were kept in dynamic conditions on an orbital shaker (IKA), rotating at 80 rpm until the collection day. From day 8 of organoid culture organoids were kept in N2B27 differentiation media which only differs from the pattering media by lack of purmorphamine. Media changes were done every 3-4 days for embedded and non-embedded organoids until the day of sample collection – day 30 or day 60 of organoid culture. For organoid cultures with reduced insulin concentration in media, commercial N2 supplement was substituted by self-made N2 supplement w/o insulin starting from day 2 of organoid culture. Self-made N2 was prepared as described in Harrison et al. (32), adding human Apo-transferrin (Sigma, cat.no.T1147), Putrescine dihydrochloride powder, cell-culture tested (Sigma, cat. no. P5780), Sodium selenite powder, cell-culture tested (Sigma, cat. no. S5261), Progesterone powder, cell-culture tested (Sigma, cat. no. P8783) in cell media in the same concentrations as in the commercial N2 supplement.

All cell cultures were tested for mycoplasma contamination once per month using LookOut® Mycoplasma PCR Detection Kit (Sigma, cat.no. MP0035-1KT).

### Insulin ELISA

Insulin in fresh cell culture media and organoid supernatant was measured with Insulin ELISA (Mercodia, cat.no. 10-1113-01) following the manufacturer’s instructions. The supernatant was collected after 2 hours of incubation with cells. Before the essay media and supernatant were diluted in DMEM-F12 (Thermo Fisher Scientific, cat.no 21331046) 1:500 to ensure that insulin concentration do not exceed the measurement range of the kit. DMEM-F12 was used as a negative control for the assay as it does not contain any insulin. All samples were measured in duplicates from which the average was calculated.

### Western Blot

IPSC-derived dopaminergic neurons were lysed in 1% SDS supplemented with protease inhibitors (Roche, cat no. 11697498001) and immediately boiled at 95°C for 5 minutes. Cell extracts were then sonicated at 4°C for 5 cycles (30 seconds on / 30 seconds off) with the Bioruptor Pico (Diagenode), prior to proceeding with protein quantification using the Pierce™ BCA Protein Assay Kit (Thermo Fisher Scientific, cat. no. 23225). 10µg of proteins were resolved using SDS-PAGE (4-12% Bis-Tris precast gels in buffer MES), followed by transfer on nitrocellulose membrane in Tris-Glycine buffer containing 20% ethanol (1h15 at 400mA). Membranes were blocked for 1h in TBS buffer containing 0.2% Tween (TBS-T) and 5% BSA, and then incubated overnight at 4°C with the primary antibodies (see Table S2). Membranes were then washed in TBS-T for 30 minutes, followed by 1h incubation at RT with the corresponding HRP-linked anti-rabbit or anti-mouse secondary antibodies. After additional washes in TBS-T for 30 minutes, proteins were revealed using the ECL prime and images acquired with the STELLA 8300 imaging system (Raytest).

For the analysis of midbrain organoids, eight non-embedded organoids were pooled together from three independent organoid generation rounds and lysed using RIPA buffer (Abcam, cat no. ab156034) supplemented with cOmplete^TM^ Protease Inhibitor Cocktail (Roche, cat no. 11697498001) and Phosphatase Inhibitor Cocktail Set V (Merck, cat..no. 524629) for 20 minutes on ice. To disrupt DNA, protein lysates were sonicated for 10 cycles (30 seconds on / 30 seconds off) using the Bioruptor Pico (Diagenode). Samples were centrifuged at 4°C for 30 minutes at 14,000g. Protein concentration was determined using the Pierce™ BCA Protein Assay Kit (Thermo Fisher Scientific, cat. no. 23225). The concentration of samples was adjusted by dilution with RIPA buffer. Samples were then boiled at 95°C for five minutes in denaturalizing loading buffer. 20µg of protein of each sample was used for protein detection. Proteins were separated using SDS polyacrylamide gel electrophoresis (Bolt™ 4-12% Bis-Tris Plus Gel, Thermo Fisher Scientific) and further transferred onto a PVDF membrane (Thermo Fisher Scientific) using iBlot™ 2 Gel Transfer Device (Thermo Fisher Scientific). For IRS1 protein, protein separation was achieved using NuPAGE™ 3 to 8%, Tris-Acetate gel (Thermo Fisher Scientific). After transfer, membranes were fixed in 0.4% paraformaldehyde (PFA) in PBS for 30 minutes at room temperature (RT) and then washed twice with PBS for 10 minutes. Membranes then were blocked for one hour at RT in 5% skimmed milk powder or 5% BSA (for phosphorylated proteins) and 0.2% Tween in PBS following overnight incubation at 4°C with the primary antibodies prepared in 5% skimmed milk powder or 5% BSA (for phosphorylated proteins) and 0.2% Tween in PBS. A list of primary antibodies see in Table S2. After incubation with primary antibodies, membranes were washed three times for five minutes with 0.02% Tween in PBS and incubated for one hour at RT with HRP-linked antibodies – anti-rabbit (VWR, cat.no. NA934) or anti-mouse (VWR, cat.no. NA931). Membranes were then washed three times for ten minutes with 0.02% Tween in PBS. The signal was developed using a chemiluminescent substrate (Life Technologies, cat.no. 34580) and imaged with STELLA 8300 imaging system (Raytest). After developing phospho-AKT and phosphor-FOXO1 protein signal, membranes were stripped using Restore™ Western Blot Stripping Buffer (Thermo Fisher Scientific, cat.no. 21063) for 15 minutes at 37°C, and then washed twice for 10 min with 0.2% Tween in PBS at RT. Membranes were blocked again for one hour before overnight incubation with the AKT pan or FOXO1 pan antibody respectively.

Band intensity was quantified using ImageJ software. Phosphorylated protein relative abundance was determined in comparison to the total protein. For all other proteins, relative abundance was determined in comparison to the housekeeping protein β-Actin. Results were batch normalised to the mean intensity of all samples.

### Extracellular metabolite detection with GC-MS

Spent medium pooled from five embedded organoids from each cell line and condition in triplicates were collected 72 hours after the last media change. Extracellular metabolite extraction and subsequent GC-MS analysis were performed as previously described (33).

Metabolite derivatisation was performed using a multi-purpose sampler (Gerstel). Dried polar sample extracts were dissolved in 20 μL pyridine, containing 20 mg/mL of methoxyamine hydrochloride (Sigma-Aldrich), and incubated under shaking for 120 min at 45 °C. After adding 20 μL N-methyl-N-trimethylsilyl-trifluoroacetamide (MSTFA; Macherey-Nagel), samples were incubated for an additional 30 min at 45 °C under continuous shaking.

GC-MS analysis was performed using an Agilent 7890B GC – 5977A MS instrument (Agilent Technologies). A sample volume of 1 μL was injected into a Split/Splitless inlet, operating in split mode (10:1) at 270 °C. The gas chromatograph was equipped with a 5 m guard column + 30 m (I.D. 250 μm, film 0.25 μm) DB-35MS capillary column (Agilent J&W GC Column). Helium was used as the carrier gas with a constant flow rate of 1.2 mL/min.

The GC oven temperature was held at 90 °C for 1 min and increased to 270 °C at 9 °C/min. Then, the temperature was increased to 320 °C at 25 °C/min and held for 7 min. The total run time was 30 min. The transfer line temperature was set constantly to 280 °C. The mass selective detector (MSD) was operating under electron ionisation at 70 eV. The MS source was held at 230 °C and the quadrupole at 150 °C. Mass spectrometric data were acquired in full scan mode (m/z 70 to 700).

All GC-MS chromatograms were processed using MetaboliteDetector (v3.2.20190704) (34). Compounds were annotated by retention index and mass spectrum using an in-house mass spectral library. The following deconvolution settings were applied: Peak threshold: 10; Minimum peak height: 10; Bins per scan: 10; Deconvolution width: 5 scans; No baseline adjustment; Minimum 15 peaks per spectrum; No minimum required base peak intensity. The dataset was normalized by using the response ratio of the integrated peak area of the analyte and the integrated peak area of the internal standard. Final metabolite relative abundance was normalized to the total area of organoids from each cell line and condition from the respective batch.

### Multielectrode array

Multielectrode array (MEA) recordings were performed using the Maestro system from Axion BioSystems. Firing activity was recorded for midbrain organoids at day 30 and day 60 of organoid culture from three independent organoid generations for both time points. Three days before recording non-embedded organoids were placed into CytoView MEA 48-Black (Axion, cat.no. M768-tMEA-48B-5) plate precoated with 0.1 mg/ml poly-D-lysine hydrobromide (Sigma, cat.no.P7886) for 24 hours following 10 µg/ml laminin (Sigma, cat.no.L2020) coating for another 24 hours. Organoids were positioned at the centre of the well and covered with a droplet (20µ) of geltrex (Invitrogen, cat.no.A1413302) to secure organoid attachment. Recorded firing activity was exported using AxIS software (v.2.1. Axion) and analysed with R (version 4.2.2). The organoid activity was analysed from the active electrodes. Electrodes that did not record any activity or recorded three or fewer spikes in five minutes were excluded from the analysis. Statistical analysis was determined with Wilcoxon signed-rank test. Outliers were excluded using interquartile range (IQR) 1.5 method based on 25^th^ and 75^th^ percentiles. Data was batch normalised by the average value of the batch in each time point.

### Local field-potential recording

Individual organoids were randomly selected from three different organoid generation batches and recorded between day 30 and day 33 or day 60 and day 63. For each organoid recordings were performed in multiple local fields. Each measurement is depicted as a separate data point. Live organoids were stuck on the bottom of the slice recording chamber using a customised platinum anchor and were perfused with a fast-flow rate (8-10 ml/min) warmed (37°C) extracellular solution containing (in mM): 140 NaCl, 2 CaCl2, 1 MgCl2, 4 KCl, 10 glucose, and 10 HEPES (pH 7.3 with NaOH), bubbled with an air pump at room atmospherically condition. Solution temperature was controlled by Perfusion Temperature Controller (Scientifica). To avoid bubbles in the recording chamber, a bubble trap was employed in the inlet line, perfused by gravity drip. Local field-potential (LFP) activity was recorded using a patch pipette prepared from thin-borosilicate glass (1.5mmOD, 0.84mmID, World Precision Instruments, cat.no. 1B150F-4) pooled and fire-polished on a Flaming/Brown micropipette puller (Sutter Instrument, Model P-1000) to a final resistance of 1,5-2,5 MΩ when filled with standard extracellular solution. LFP recordings were performed with a Multiclamp 700B/Digidata1550 system (MolecularDevices) in a current-clamp configuration on a visually identified zone of healthy neurons on the organoid using a slice scope upright microscope (Scientifica). The patch pipette was pushed against the organoid for a depth of 40-60 µm until a small increase in the standard deviation of the recorded voltage was seen.

Data were acquired with Clampex software (v.11.2, MolecularDevices) at 3 kHz sample frequency and filtered at 3 kHz with a low-pass Bessel filter and 1Hz with a high-pass filter. All data were analyzed offline with Clampex software (v.11.2, MolecularDevices). The detection of LFP was done in 5-minute temporary windows recording with the threshold search function that recognizes only spikes over 6X standard deviation. Bursts with a minimum of 3 spikes and an inter-spike interval of ≤150 msec were detected and considered for analysis.

### Immunofluorescence staining for NESC characterisation

For immunofluorescent characterization, NESCs were seeded on geltrex-coated coverslips placed in a 24-well cell culture plate. At 60-70% of confluency, NESCs were fixed for 15 min at RT with 4% *Paraformaldehyde* (PFA). Then cells were washed 3x for 5 min with PBS, which was followed by cell permeabilization with 0.3% Triton X-100 in PBS for 15 min at RT. Afterwards, cells were washed 3x for 5 min with PBS and then blocked with 10% fetal bovine serum (FBS) in PBS for 1h at RT. Incubation with primary antibodies (Table S3) was done overnight at 4°C. Antibodies were diluted in 3% FBS in PBS. After overnight incubation, cells were washed 3x for 5 min with PBS and incubated for 1h at RT with Alexa Fluor^®^ conjugated secondary antibodies (Table S3) and nuclei stain Hoechst 33342 (Invitrogen cat.no. 62249). After 3x washes for 5 min with PBS, coverslips were mounted on a glass slide for image acquisition.

### Immunofluorescence staining of organoid sections

Organoids were fixed with 4% PFA for six hours at RT and then washed three times with PBS for 15 minutes. Individual organoids were embedded in 3% low-melting point agarose (Biozym Scientific GmbH, cat.no. 840100). Using a vibrating blade microtome (Leica VT1000s), organoids were sliced into 70 µm sections. Sections were permeabilized with 0.5% Triton X-100 for 30 minutes at RT and then blocked with blocking buffer (2.5% normal donkey serum, 2.5 % BSA, 0.01% Triton X-100 and 0.1 % sodium azide). Sections were incubated with primary antibodies diluted in a blocking buffer for 48 hours at 4 °C on an orbital shaker. Primary antibodies are listed in Table S3. After incubation with primary antibodies, sections were washed three times for 5 minutes with 0.01% Triton X-100 in PBS. Then sections were incubated with nuclei stain Hoechst 33342 (Invitrogen cat.no.62249) in dilution of 1:10 000 and The Alexa Fluor^®^ conjugated secondary antibodies (Table S3) for two hours at RT on an orbital shaker. Sections were washed twice for 5 minutes with 0.01% Triton X-100 in PBS and once with MiliQ. Sections were mounted on slides (De Beer Medicals, cat.no.BM-9244) and covered with mounting media Fluoromount-G® (SouthernBiotech, cat.no.0100-01).

Apoptotic cells were stained using Tunel In Situ Cell Death Detection Kit, Fluorescein (Merck, cat.no. 11684795910). Kit components were integrated into the staining procedure. Staining with primary antibodies was done as described above. Secondary antibodies and Hoechst were diluted in the Tunel kit buffer. Sections were incubated for one hour at RT on an orbital shaker following washing and mounting steps as described above.

### Image acquisition and analysis

High-content imaging was performed using the Yokogawa CV8000 high-content screening microscope with a 20x objective. One to three sections from at least three independent organoid generation rounds were analyzed per cell line and condition for each staining combination. Acquired images were analyzed with a customized pipeline using Matlab (v.2021a) as described in Bolognin et al, 2018 (35) and Monel et al, 2020 (36). Sections with non-optimal mask for a target protein or out of focus were excluded from further analysis. Data was batch normalised by the average value of the batch per feature and per time point. Outliers were excluded using interquartile range (IQR) 1.5 method based on 25^th^ and 75^th^ percentiles. Qualitative images were acquired using a confocal laser scanning microscope (Zeiss LSM 710) with either a 20x, 40x or 60x objective and processed with Zen Software (Zeiss)

### RNA sequencing and analysis

The total RNA was extracted from 12 embedded organoids, which were pooled from three independent organoid generations (four organoids from each generation per cell line and per condition). RNA isolation was achieved using Trizol reagent (Thermo Fisher Scientific, cat. no. 15596026) following the manufacturer’s protocol. Library preparation was done using TruSeq stranded mRNA library preparation kit (cat. no. 20020594, Illumina) and sequenced on NextSeq2000.The raw RNA-seq data were pre-processed using the software package “Rsubread” (version 1.34.7) (37).

### Metabolic modelling

Gene expression from RNA sequencing was normalized to the transcripts per kilobase million (TPM) reads. The Median of TPM expression was calculated between the three cell lines per IR and IS condition to enable the generation of one context-specific model per condition. Gene EntrezID as required input in the modelling pipeline was mapped using the clusterProfiler package (version 3.9) (42) in R (version 3.6.2). Context-specific models were reconstructed from the stoichiometrically and flux-consistent subset of the human genome-scale metabolic network Recon3 (43) using the rFASTCORMICS pipeline (44). Additionally, models were constrained by the cell culture media composition, allowing uptake of only those metabolites present in the media. Models were analyzed in Matlab (v.2019b) using the COBRA toolbox (45).

Models were compared by reaction composition, using the setdiff function in Matlab. Reactions that were unique for one of the models, were assigned to the subsystem. Subsystems with more than eight reactions unique for one of the models were selected for further analysis. For selected subsystems, the number of unique reactions was expressed as a percentage of the total number of reactions in the respective subsystem in Recon.

Flux of the ATP demand/generation ‘DM_atp_c_ ‘reaction in each model was estimated by the addition of qualitative constraints of glucose uptake ‘EX_glc_D[e]’, lactate secretion ‘EX_lac_L[e]’ and biomass reaction. Biomass increase per hour was calculated based on embedded organoid area increase between day 16 and day 30 of organoid culture. Dry weight (DW) for a single cell was calculated considering that the total protein fraction within the dopaminergic neurons neurons is 55.93%(46). Glucose uptake and lactate secretion were calculated as mmol/gDW*h. The concentration change of glucose and lactate within 72 hours for DW was estimated by comparing the relative abundance determined with GC-MS to metabolite concentration in the commercial media. Glucose and lactate relative abundance for each model was calculated as a median between the samples, after averaging values between technical replicates. The lower and upper bound for glucose uptake and lactate secretion was set as +/- 10% of the calculated flux, while the lower and upper bound for biomass were kept exact. Next, estimated ATP demand for each model was used as an objective function set as the lower bound for the ‘DM_atp_c_’ reaction. Further, FVA using IBM_CPLEX solver (version 12.10) was performed for shared reactions between the two models for the subsystems of interest – all subsystems determined to have the largest differences in reaction composition with the addition of glycolysis, TCA and OXPHOS. An average of min and max fluxes was calculated for visualization of flux variability between the two models. ATP perturbations were performed by setting the upper bound of reactions of interest to 0 and re-estimating ATP demand flux.

### Intracellular ATP assay

Intracellular ATP was measured using luminescence-based CellTiter-Glo® 3D Cell Viability Assay (Promega, cat.no. G9681) following the manufacturer’s instructions. Three organoids per cell line and per condition were transferred into the imaging plate (PerkinElmer, cat.no.6055300)– one organoid per well. Media leftovers were aspirated and 50 µl of CellTiter-Glo® reagent was added to each well. The plate was incubated for 30min on a shaker at RT. Luminescence was measured using a Cytation5 M cell imaging reader. The experiment was repeated for five independent organoid generation rounds at day 30 and day 60 of organoid generation. The luminescence readout per sample of each batch was normalized to the mean luminescence of the batch. A median of three organoids of each cell line and conditions was calculated and normalized to the size factor. The size factor for each cell line and condition was determined as a mean of random 20 organoids at day 30 and 10 organoids at day 60 from five independent organoid generation rounds. The area of organoids was measured using the Cytation5M cell imaging reader.

### Fatty acid dependency assay (Seahorse XF Mito fuel flex test)

The rate of fatty acids oxidation was determined in 30 days old midbrain organoids following Seahorse XF Mito fuel flex test protocol (Agilent) and using Seahorse XFe96 Spheroid FluxPak (Agilent, cat.no. 102905-100). The utility plate was coated with Corning® Cell-Tak™ Cell and Tissue Adhesive (Corning, cat.no. 354241) and for one hour incubated at 37^0^C before organoid seeding in the plate. Organoids were incubated for one hour in Seahorse XF DMEM medium, pH 7.4 (Agilent, cat.n.103575-100) supplemented with 1 mM pyruvate, 2 mM glutamine, and 10 mM glucose at 37^0^C in the non-CO2 incubator. Assay was performed for five independent organoid generation rounds, using 6-8 organoids per cell line and condition in each assay run. Results were normalised to the size factor. The size factor for day 30-old organoids for each cell line and condition was determined as a mean of random 20 organoids from five independent organoid generation rounds. The area of organoids was measured using the Cytation5M cell imaging reader. Data were analyzed with the Seahorse Wave Desktop software (Agilent). Wells with aberrant OCR patterns were excluded from each of the individual experiments.

### Stable isotope tracing sample preparation

Cell culture media was prepared in DMEM w/o glucose (Themor Fisher Scientific, cat.no. A1443001) and Neuobasal A w/o glucose (Thermo Fisher Scientific, cat.no. A2477501). [U-^13^C] glucose stable isotope tracer (Eurisotop, cat.no.CLM-1396) was added to the media to a final concentration of 21.25mM. 30 days old, non-embedded organoids were incubated in the tracer medium for 24h at 37°C, 5 % CO_2_ in a humidified incubator. Eight organoids per cell line and condition were pooled to perform metabolite extraction. Collected samples were first washed with 1xDPBS and then resuspended in 80μl ice-cold extraction solvent (50% MeOH, 30% ACN, 20% H2O). Samples were incubated in a thermo shaker at 4°C, 300 rpm for 5 min and then centrifuged at full speed at 4°C for 10 min. 70μl of supernatant was transferred into LC-MS vials and stored at −80°C until the analysis.

### LC-MS Measurement

Metabolite analysis was performed using a Vanquish® Flex UHPLC system (Thermo Scientific) coupled to a Q –Exactive® Plus Orbitrap® mass spectrometer (Thermo Scientific). Chromatography was carried out with a SeQuant® ZIC®-pHILIC 5µm polymer 150 x 2.1 mm column (Merck AG) protected by a SeQuant® ZIC®-pHILIC Guard 20 x 2.1 mm pre-column. The temperature of the column was kept at 45°C and the flow rate was set to 0.2 ml/min. The mobile phases consisted of 20 mmol/L ammonium carbonate in water, pH 9.2 and supplemented with 5µM of medronic acid (Eluent A) and Acetonitrile (Eluent B), loading and gradient occurs at 0,2mL/min flow rate; loading step last 2min with 80% B; then starts a linear gradient ranging from 80% B to 20% B in 15 min; at 18 min starts the regeneration step with 80% B; from 20 min until 24,5 min column is re-equilibrated with 80% B at flow rate of 0.4mL/min. 5 µL of sample were injected into the machine. The MS experiment was performed using electrospray ionization with polarity switching enabled. Source parameters were applied as written here: sheath gas flow rate, 30; aux gas flow rate, 10; sweep gas flow rate, 0; spray voltage, 4 kV (+) / 3.5 kV (-); capillary temperature, 350°C; S-lense RF level, 50; aux gas heater temperature, 100°C. The Orbitrap mass analyzer was operated at a resolving power of 70,000 (at 200 m/z) in full-scan mode (scan range: *m/z* 75-1000; automatic gain control target set to 1e6 charges with a maximum injection time: 250 ms). Data from LC-MS measurements were acquired with the Thermo Xcalibur software (v.4.3.73.11) and analyzed using TraceFinder (v.4.1). Natural isotope abudance substraction and subsequent analysis was performed as previously described using in-house scripts (51).

### Reactive oxygen species detection with fluorescence-activated cell sorting (FACS)

To quantitatively determine cytoplasmic reactive oxygen species (ROS) levels in live cells, 4-5 non-embedded organoids at day 30 of organoid culture were pooled per sample and dissociated using Accuatse (Sigma, cat.no. A6964). Samples were first incubated with Accutase for one hour at 37^0^C under dynamic conditions, following trituration with a 1000 or 200 μl pipette (depending on organoid size) to support organoid dissociation into single cells. Reducing incubation time to 30 min, these steps were repeated until organoids were fully dissociated. Each sample then was split into two 2ml Eppendorf tubes as technical replicates for the assay. Samples were centrifuged for 3 min at 500 g and Accutase was replaced with 500ul assay medium containing 2μM 2’,7’ –dichlorofluorescein diacetate (DCFDA) (Thermo Fisher Scientific, cat.no. D399) and 1:1000 live-dead stain Zombie NIR (Biolegend, cat.no. 423106) in DPBS (Thermo Fisher Scientific, cat.no. 14190250). Cells were incubated with the assay media containing the fluorescent dyes for 20 min at 37^0^C. Samples were then run on BD LSRFortessa flow cytometer, acquiring 10,000 single-cell events (DCFDA+, live cells) per sample. Fluorescence intensity was analysed with FlowJo software (v.10.7.2). Data of two technical replicates per sample were averaged and the percentage of positive cells to the DCFDA staining was batch normalised to the mean percentage of the batch.

### FACS-based glucose uptake assay using a 2-NBDG probe

Glucose uptake was measured using fluorescently tagged, non-metabolizable glucose analogue 2-(N-(7-nitrobenz-2-oxa-1,3-diazol-4-yl) amino)-2-deoxyglucose (2-NBDG) (Thermo Fisher Scientific, cat.no. 11569116). Samples were prepared as described in the section: *Reactive oxygen species detection with fluorescence-activated cell sorting (FACS).* Cells were incubated for 30 min at 37^0^C with 1:500 2-NBDG and 1:1000 live-dead stain Zombie NIR (Biolegend, cat.no. 423106) in cell culture media with either high or low insulin concentration. Samples were run on BD LSRFortessa flow cytometer, acquiring 10,000 single-cell events (2-NBDG+, live cells) per sample. Fluorescence intensity was analysed with FlowJo software (v.10.7.2). Data of two technical replicates per sample were averaged and the fluorescence intensity was batch normalised to the IR condition.

### Lipidomics

Lipidomics analysis was performed by Lipometrix (http://www.lipometrix.be) using HILIC LC-MS/MS. Samples were prepared as a pool of eight non-embedded organoids per cell and per condition from four independent organoid generation rounds. Samples were collected on day 30 of organoid culture, washed once with ice-cold PBS, snap-frozen with liquid nitrogen and stored at −80°C until the analysis. The concentration of detected lipid species was normalized to DNA content.

### Dopamine ELISA

Extracellular dopamine concentration was measured with Dopamine ELISA (Immusmol, cat no. BA-E-5300R) following the manufacturer’s instructions. Dopamine ELISA was performed using a 750 µl volume of a sample. Each sample was a pool of 400 µl of supernatant from three embedded midbrain organoids, collected at day 60 of organoid culture, 72 hours after the last media change. Samples were collected from three independent organoid generation rounds. Final dopamine concentration (nmol/l) was normalized to the mean nuclei count determined during the high-content imaging workflow from randomly selected three organoid sections per cell line and condition. Normalized results were batch-corrected against WT organoid mean values.

### Data analysis and statistics

If not stated otherwise, data were analysed and visualized in GraphPad Prism (v.9.5.1) or R software (v.4.2.2). All data were batch normalized for every experiment. Data normality was determined with the Shapiro-Wilk normality test. Comparing healthy donor midbrain organoid response to different insulin conditions, the statistical significance of non-normally distributed data was tested with a non-parametric, two-tailed Wilcoxon matched-pairs signed rank test. For normally distributed data, statistical significance was tested with a two-tailed, paired t-test. Significance asterisks represent *P*<0.05 *, *P* <0.01 **, *P* <0.001 ***. Error bars represent mean + SD.

### Ethics approval

The work with iPSCs has been approved by the Ethics Review Panel (ERP) of the University of Luxembourg and the national Luxembourgish Research Ethics Committee (CNER, Comité National d’Ethique de Recherche). CNER No. 201901/01; ivPD

## Acknowledgements

This work was mainly supported by the internal flagship project at the Luxembourg Centre for Systems Biomedicine and by the Luxembourg National Research Fund CORE grant to GA (C21/BM/15850547/PINK1-DiaPDs). Further, we acknowledge support from the National Centre of Excellence in Research on Parkinson’s Disease (NCER-PD) which is funded by the Luxembourg National Research Fund (FNR/NCER13/BM/11264123). We thank Dr Sarah Nickels and Isabel Rosety for the characterisation of iPSCs and the derivation of NESCs. Also, we would like to thank Prof. Dr. Rudi Balling for the valuable ideas, which inspired this project.

## Author contributions

A.Z. designed experiments, analysed and interpreted data, prepared figures, and wrote the original draft. J.K. analysed metabolic models. C.A. and A.V. carried out Western Blot experiments. D.F. carried out the mounting of stained organoid sections. D.F. performed local field potential recordings and data analysis. G.A. and G.G.G. designed experiments, reviewed, and edited the manuscript. C.J. performed a GC-MS metabolomics experiment. L.N. and M.J. supervised the C13 glucose tracing experiment and performed data processing and analysis. P.A. supervised high-content imaging workflow. R.H. performed RNA sequencing. E.G. performed RNA sequencing data processing and analysis. E.S. supervised the metabolic modelling part, and reviewed, and edited the manuscript. J.C.S. conceived and supervised the project and edited the manuscript.

## Rights retention statement

“This research was funded in whole by the FNR-Luxembourg. For the purpose of Open Access, the author has applied a CC BY public copyright license to any Author Accepted Manuscript (AAM) version arising from this submission.”

