## Supplementary Information for "Insulin resistance compromises midbrain organoid neural activity and metabolic efficiency predisposing to Parkinson’s disease pathology"

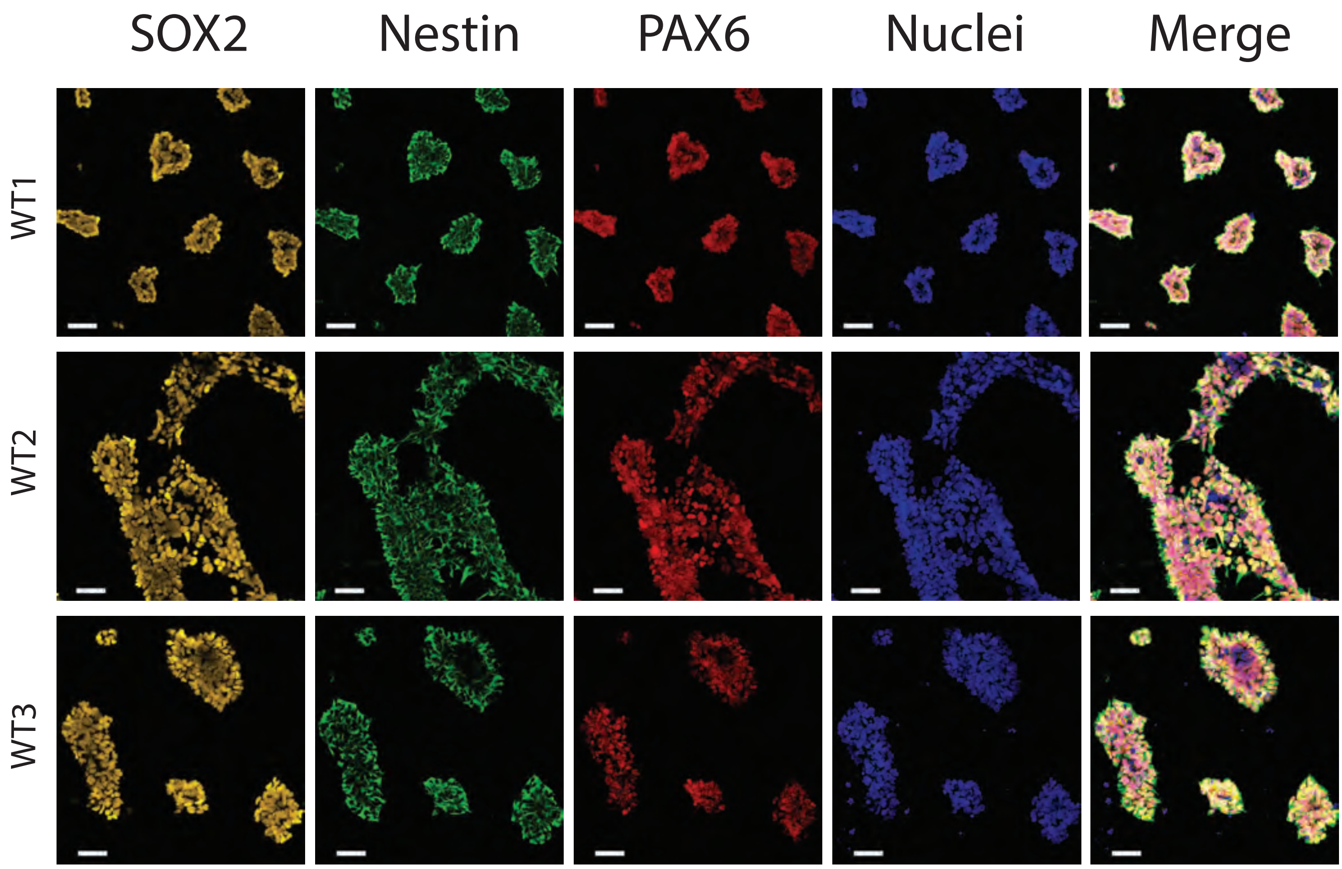


**Figure S1. NESC characterisation.** Immunofluorescent images of neuronal stem cell typical markers-SOX2 and Nestin, and neuroectoderm marker PAX6. Scale bars 50µm.

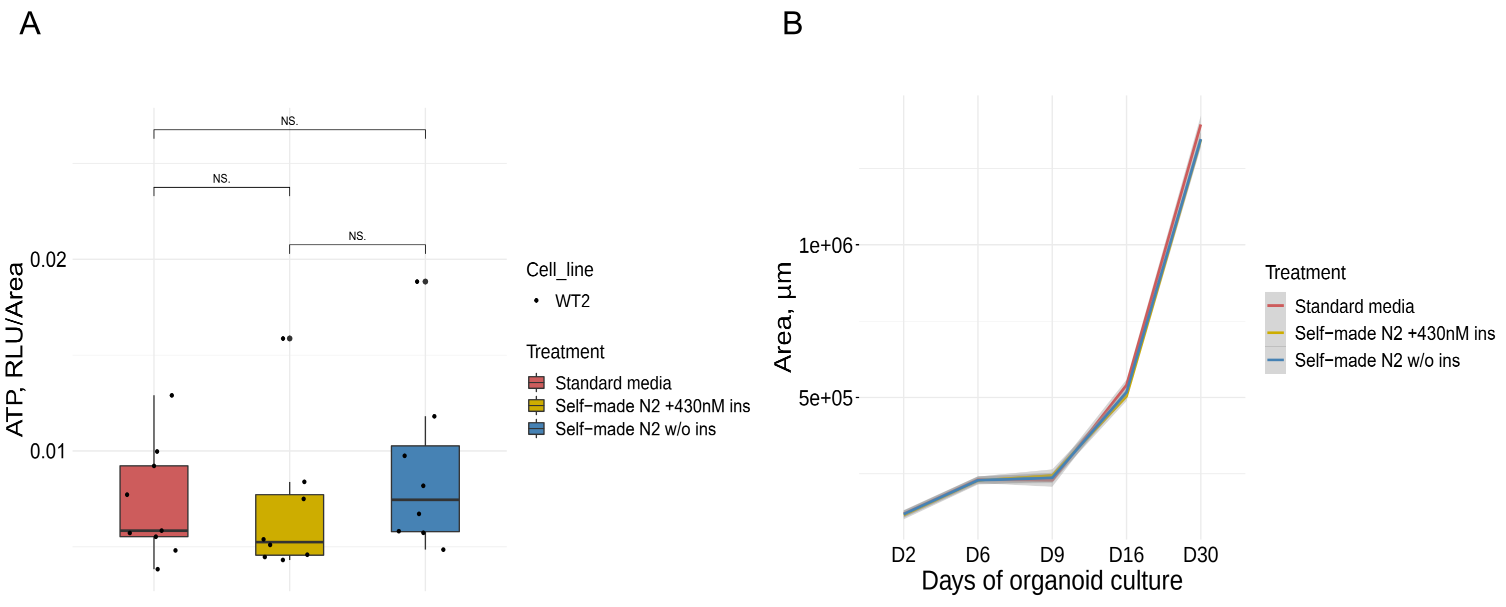


**Figure S2. Self-made N2 supplement validation in midbrain organoid culture.** Midbrain organoids cultured until day 30 in standard media (high insulin), self-made N2 media + 430nM insulin (to mimic theorethical insulin concentration in standard media), and self-made N2 media w/o insulin. (A) Intracellular ATP measurement at day 30 representing organoid viability. (B) Area measurement of midbrain organoids over the differentiation time.

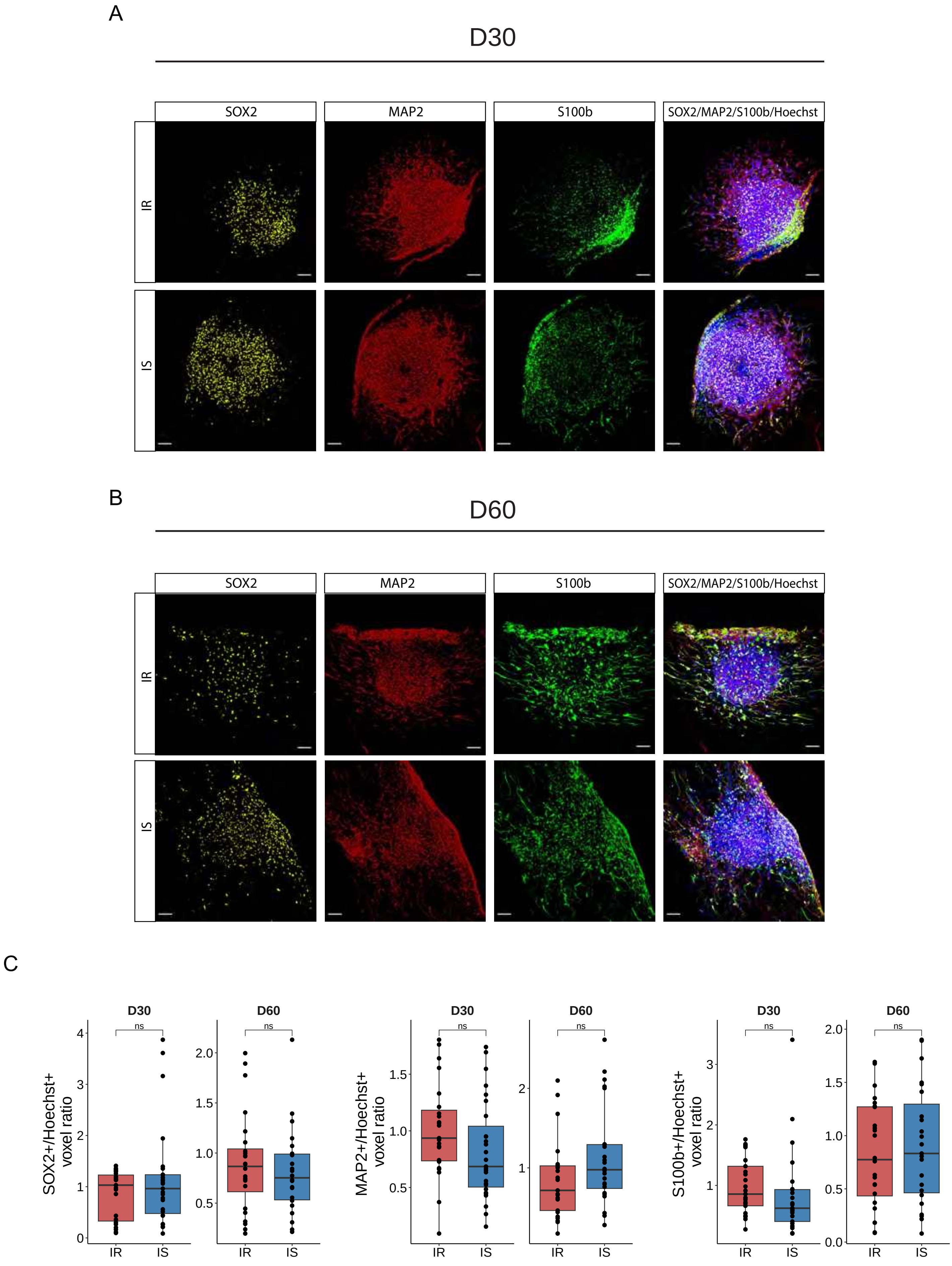


**Figure S3. Cell types in midbrain organoids.**
Representative immunofluorescent images of SOX2, MAP2 and S100b staining in insulin resistant (IR) and insulin sensitive (IS) midbrain organoids at (A) day D30 and (B) day 60. Scale bars: 100µm. (C) Quantification of SOX2, MAP2 and S100b positive area over Nuclei (Hoechst) area.

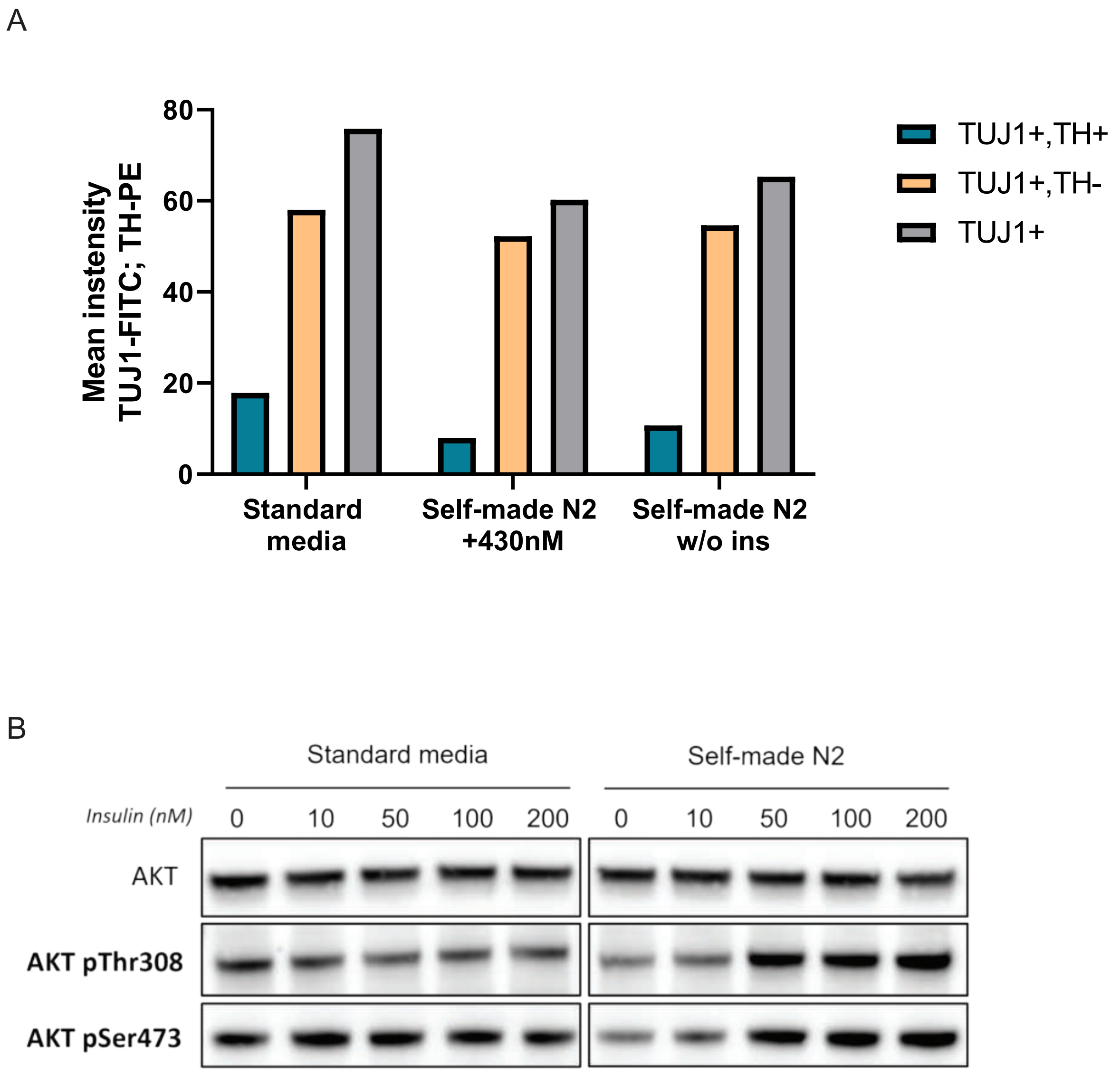


**Figure S4. Self-made N2 supplement validation in 2D iPSC-derived dopaminergic neurons.** Dopaminergic neurons were cultured in Standard media, self-made N2 media + 430nM insulin (to mimic theoretical insulin concentration in standard media), and self-made N2 media w/o insulin until day 30 of differentiation. (A) Percentage of cells quantified by FACS, expressing the indicated neuronal differentiation markers at day 30 of differentiation. (B) Activation of AKT signaling pathway was evaluated after 30 days of NESC differentiation in the two media (standard vs self-made N2), upon acute stimulation with increasing insulin doses (from 10 to 200nM) for 30 minutes. AKT phosphorylation at both T308 and S473 residues was assessed by immunoblotting and normalized against total AKT levels.

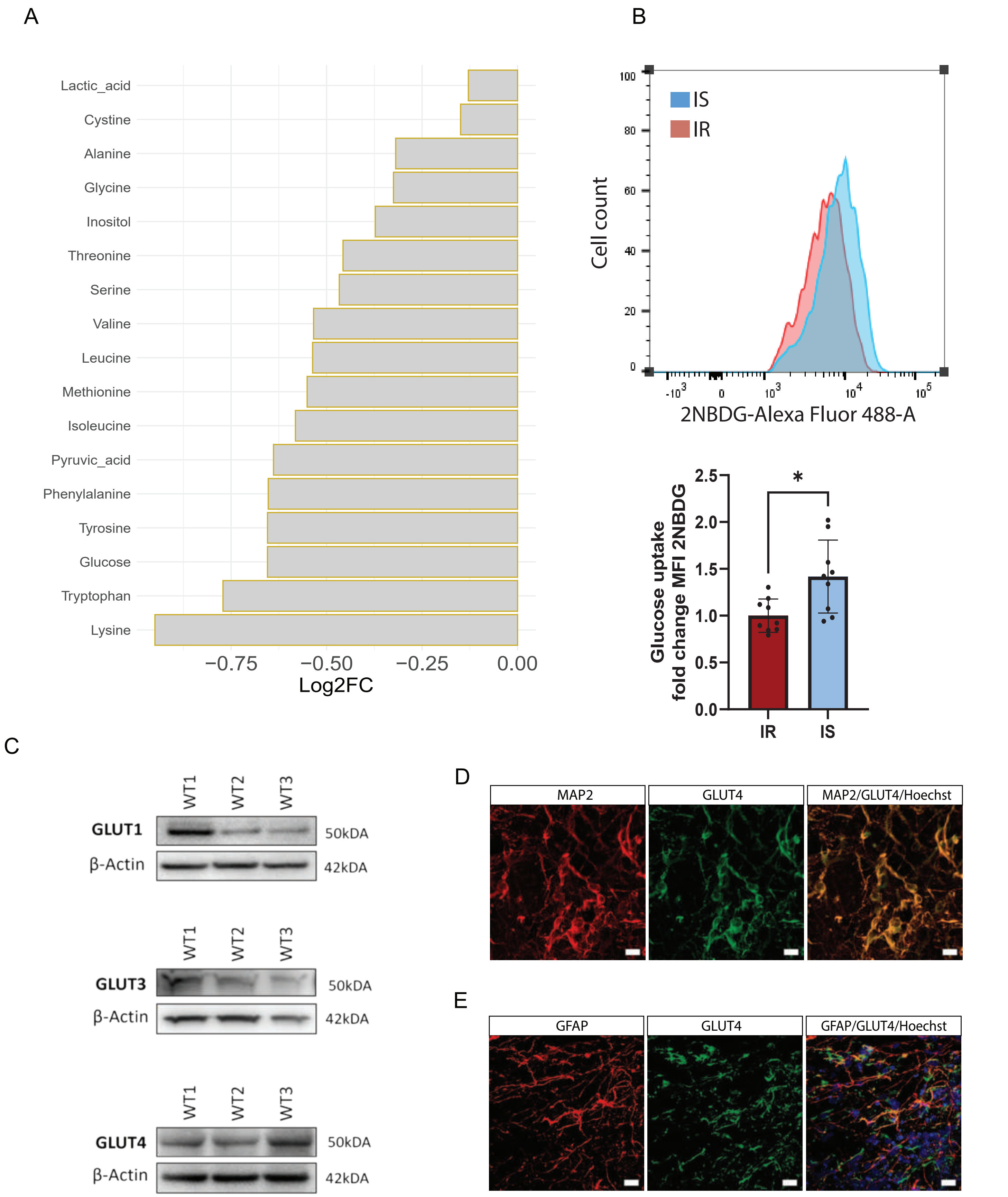


**Figure S5. Insulin-dependent metabolic changes and glucose transporter presence in midbrain organoids.** (A) GC-MS detected metabolite relative concentration in IS organoid spent media demonstrated as log2 fold change (FC) against IR samples. Metabolite relative abundance normalised to the organoid area. (B) Representative histogram of cells positive to the Alexa-Fluor 488 signal demonstrating uptake of fluorescent glucose analog - 2NBDG. Measurement performed after 30 min incubation of high (standard media) or reduced (self-made N2 media) insulin concentration after organoid dissociation. Statistical significance tested with paired t-test. (C) Representative images of Western Blots showing GLUT1, GLUT3 and GLUT4 presence in midbrain organoids. d) Representative immunofluorescent staining of MAP2 and GLUT4 at day 30 of organoid culture. Scale bars:20μm. e) Representative immunofluorescent staining of GFAP and GLUT4 at day 60 of organoid culture. Scale bars:20μm.

A

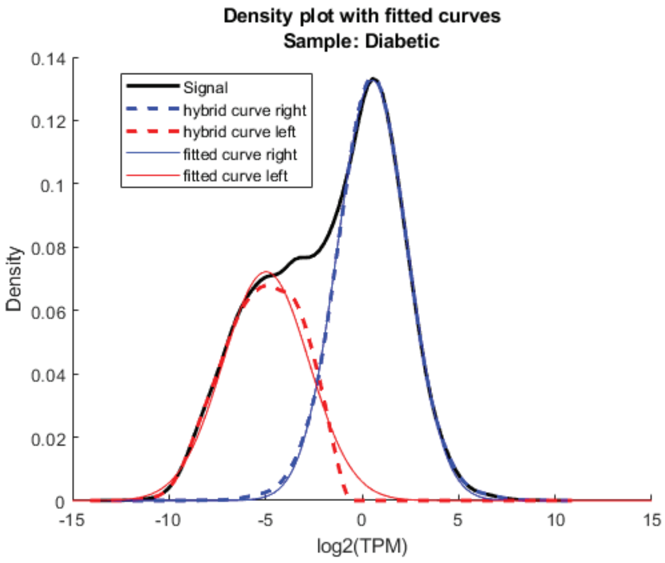


B

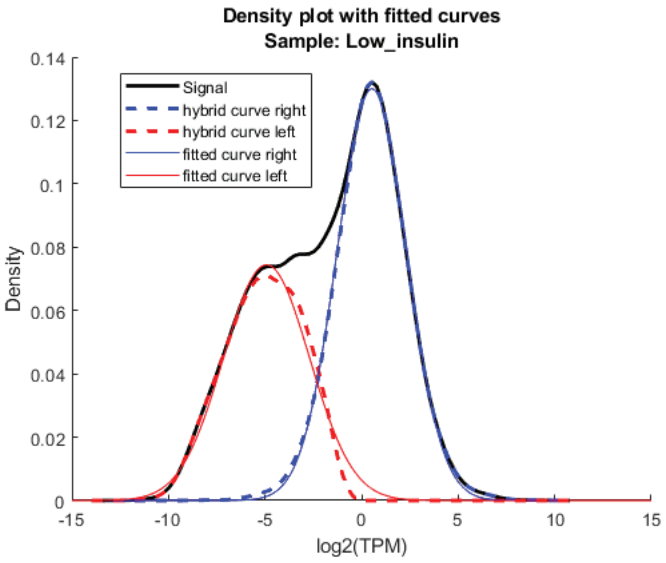


C

| Model | Genes | Rxns | Mets |
|---|---|---|---|
| IR | 1446 | 3620 | 2632 |
| IS | 1430 | 3470 | 2521 |

**Figure S6. Generation of context-specific metabolic models.** Representative histogram of gene discretization based on their expression levels of pooled data of (A) insulin resistance (IR) samples and (B) insulin sensitive S samples. (C) Generated model composition regarding the number of genes, reactions (Rxns) and metabolites (Mets) in each model.

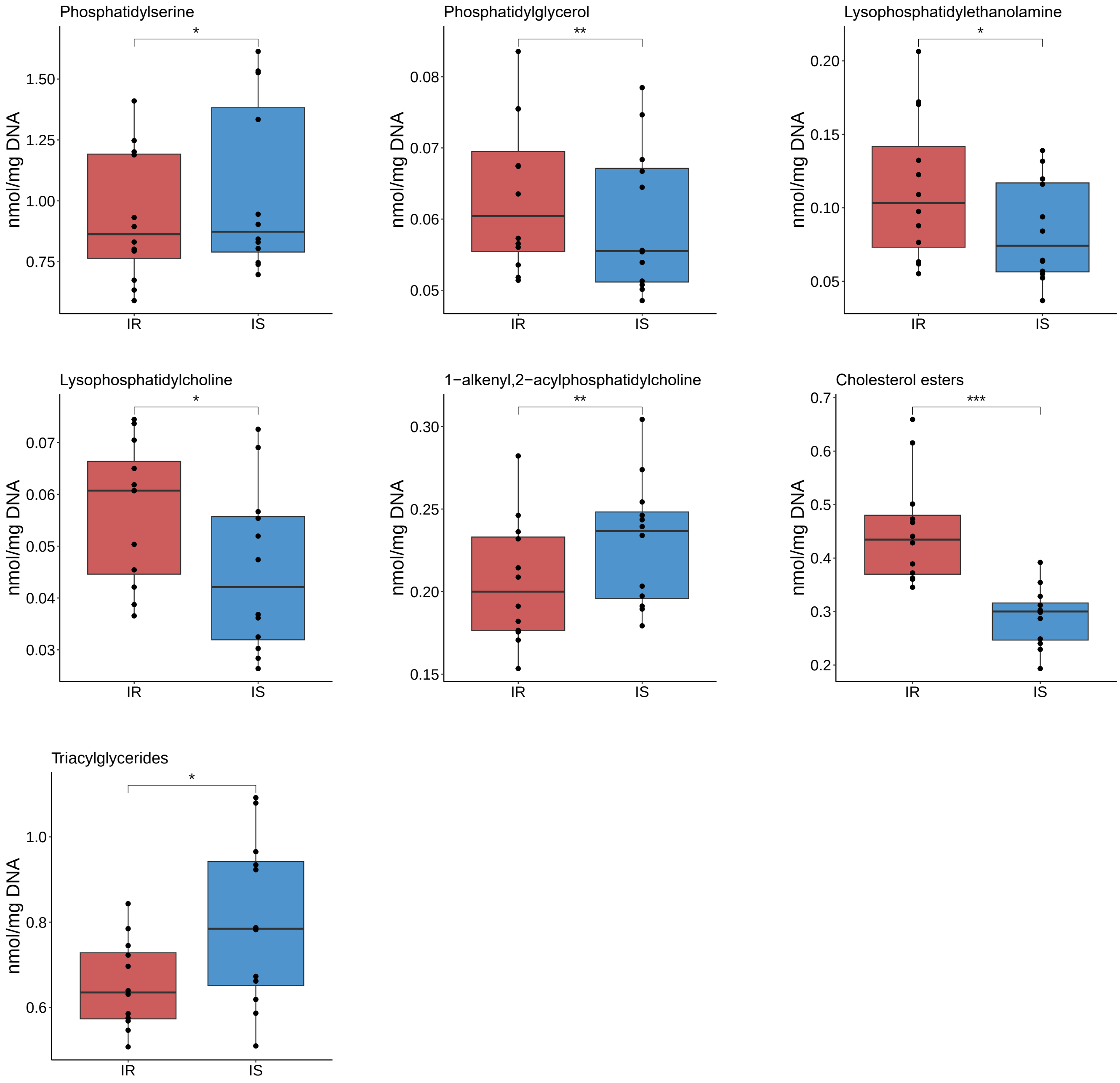


**Figure S7. Significantly differentially abundant lipid classes.**
Lipids measured by LC-MS. Final concentration normalised to the DNA content of each samples. Statistical significance tested with two-sided, paired Wilcoxon test (P<0.05 *, P <0.01 **, P <0.001 ***).

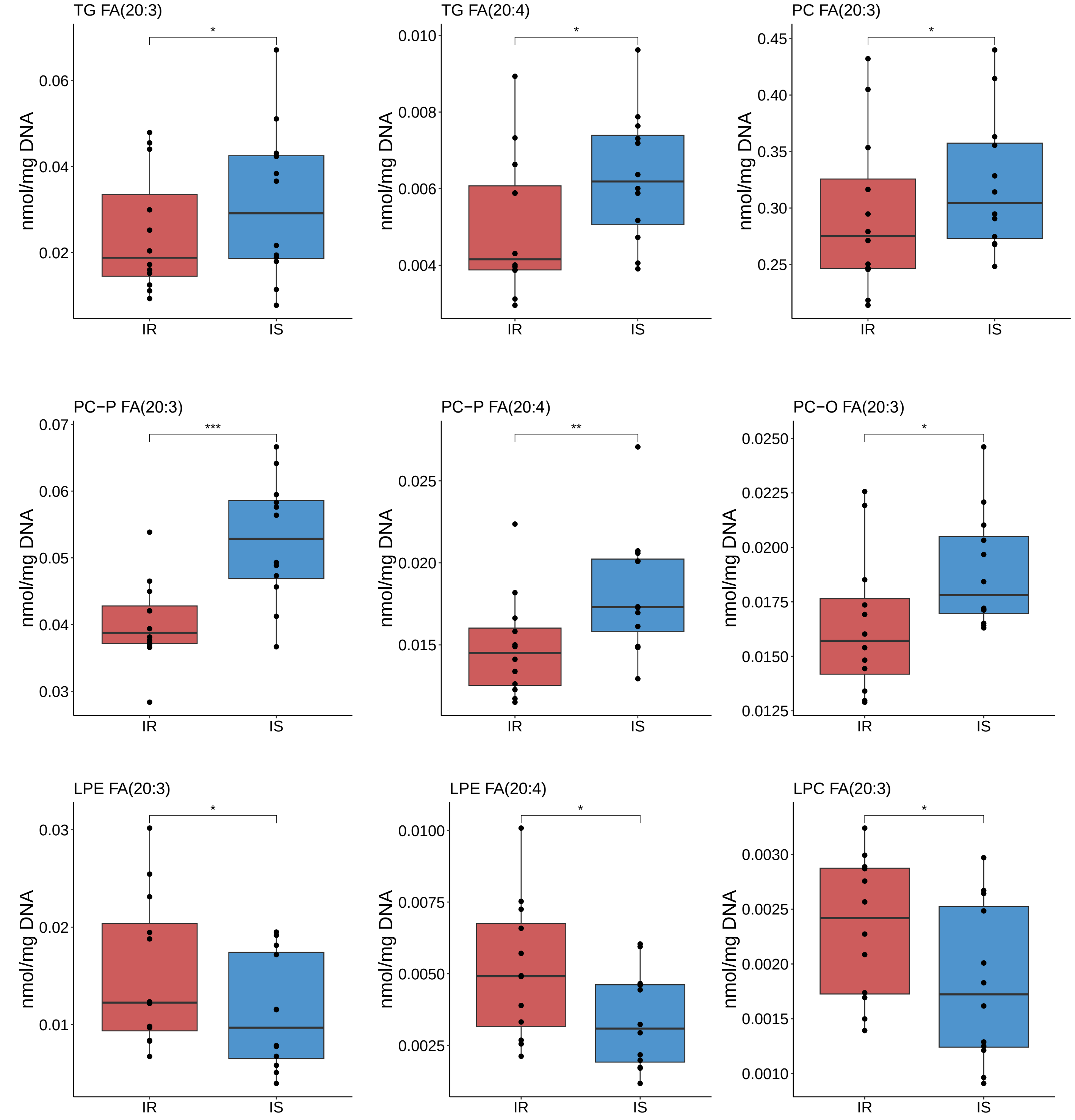


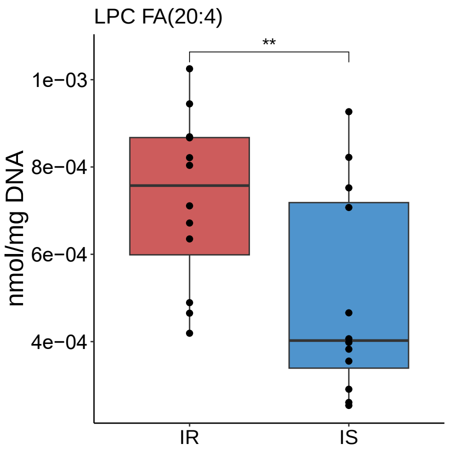


**Figure S8. Significantly differentially abundant lipid species containing 20 carbons.** Lipid species across all lipid classes with 20 carbon fatty acid in the structure with three (20:3) or four (20:4) double bonds. Final concentration normalised to the DNA content of each samples. Statistical significance tested with two-sided, paired Wilcoxon test (P<0.05 *, P <0.01 **, P <0.001 ***).

**Table S1. Cell lines used in this study**

| Sample ID | Diagnosis | Genotype | Age of sampling | Internal iPSC ID | Source of iPSC |
|---|---|---|---|---|---|
| WT1 | Healthy | wt/wt | 68 | 2.0.0.71.1.0 | IBBL / Max Planck Institute |
| WT2 | Healthy | wt/wt | 68 | 2.0.0.77.0.0 | STEMBANCC |
| WT3 | Healthy | wt/wt | 55 | 2.0.0.27.0.0 | Coriell |

**Table S2. Primary antibodies used for WB**

| Antibody | Source | Cat.no. | RRID | Species | Dilution |
|---|---|---|---|---|---|
| Phospho-Akt (Ser473) | Cell Signaling | 4060T | *AB_2315049* | Rabbit | 1:300 (organoids) 1:1000 (dopaminergic neurons) |
| Phospho-Akt (Thr308) | Cell Signaling | 13038T | *AB_2629447* | Rabbit | 1:1000 (dopaminergic neurons) |
| AKT pan | Cell Signaling | 4691T | *AB_915783* | Rabbit | 1:500 (organoids) 1:2000 (dopaminergic neurons) |
| IRS1 | Cell Signaling | 2382 | *AB_330333* | Rabbit | 1:300 |
| GLUT1 | Proteintech | 21829-1-AP | *AB_10837075* | Rabbit | 1:300 |
| GLUT3 | Proteintech | 20403-1-AP | *AB_10694437* | Rabbit | 1:300 |
| GLUT4 | Proteintech | 66846-1-Ig | *AB_2882186* | Mouse | 1:300 |
| βeta-Actin | Cell Signaling | 3700S | *AB_2242334* | Mouse | 1:10 000 |

**Table S3. Primary and secondary antibodies used for immunofluorescence stainings**

| Antibody | Source | Cat.no. | RRID | Species | Dilution |
|---|---|---|---|---|---|
| Nestin | BD Bioscience | 611659 | *AB_399177* | Mouse | 1:200 |
| PAX6 | Biolegend | 901302 | *AB_2749901* | Rabbit | 1:300 |
| TH | Abcam | ab112 | *AB_297840* | Rabbit | 1:1000 |
| TH | Sigma | T2928 | *AB_477569* | Mouse | 1:200 |
| MAP2 | Abcam | ab92434 | *AB_2138147* | Chicken | 1:1000 |
| GLUT4 | Proteintech | 66846-1-Ig | *AB_2882186* | Mouse | 1:500 |
| GFAP | Millipore | AB5541 | *AB_177521* | Chicken | 1:1000 |
| S100b | Sigma | S2532 | *AB_477499* | Mouse | 1:500 |
| SOX2 | R&D Systems | AF2018 | *AB_355110* | Goat | 1:200 |
| Anti-chicken 647 | Jackson Immuno Research | 703-605-155 | *AB_2340379* | Donkey | 1:1000 |
| Anti-rabbit 488 | Invitrogen | A21206 | *AB_2535792* | Donkey | 1:1000 |
| Anti-rabbit 568 | Invitrogen | A10042 | *AB_2534017* | Donkey | 1:1000 |
| Anti-goat 568 | Invitrogen | A11057 | *AB_2534104* | Donkey | 1:1000 |
| Anti-mouse 488 | Invitrogen | A21202 | *AB_141607* | Donkey | 1:1000 |
| Anti-mouse 568 | Invitrogen | A10037 | *AB_2534013* | Donkey | 1:1000 |
| Anti-goat 647 | Invitrogen | A21447 | *AB_2535864* | Donkey | 1:1000 |
